# Two-factor synaptic consolidation reconciles robust memory with pruning and homeostatic scaling

**DOI:** 10.1101/2024.07.23.604787

**Authors:** Georgios Iatropoulos, Wulfram Gerstner, Johanni Brea

**Author notes:** Contributing authors.

## Abstract

Memory consolidation involves a process of engram reorganization and stabilization that is thought to occur primarily during sleep through a combination of neural replay, homeostatic plasticity, synaptic maturation, and pruning. From a computational perspective, however, this process remains puzzling, as it is unclear how the underlying mechanisms can be incorporated into a common mathematical model of learning and memory. Here, we propose a solution by deriving a consolidation model that uses replay and two-factor synapses to store memories in recurrent neural networks with sparse connectivity and maximal noise robustness. The model offers a unified account of experimental observations of consolidation, such as multiplicative homeostatic scaling, task-driven synaptic pruning, increased neural stimulus selectivity, and preferential strengthening of weak memories. The model further predicts that intrinsic synaptic noise scales sublinearly with synaptic strength; this is supported by a meta-analysis of published synaptic imaging datasets.

## 1. Introduction

The ability to store and retrieve remote memory is thought to rely on a distributed network of neurons located primarily in the cortical areas of the brain [1–4]. This view is supported by anatomical studies, showing that cortical circuits are highly recurrent and, thus, particularly conducive to information storage [5–7]. In an effort to unify these findings, models of long-term memory are today often based on the concept of attractor networks [8]. The basic idea of this approach is to represent local cortical circuits with a recurrent neural network, in which each memory corresponds to a distinct pattern of activity that acts as an attractor of the network’s dynamics [9, 10]. In this context, memory encoding is modeled by configuring the connections of the network to imprint activity patterns as stable attractors. When this is done optimally, memory storage is saturated and the network reaches critical capacity [11, 12]. This state is particularly significant. In a series of recent studies, attractor networks operating close to critical capacity have been shown to mimic several dynamical and structural motifs observed in cortical circuits, thereby suggesting that optimal storage is an organizing principle of cortical connectivity [13–16]. However, it is unclear how such optimality can emerge in biology, and the precise role of synaptic plasticity in this process remains unknown.

In the experimental literature, the process whereby memories are stabilized and reshaped for long-term storage is generally referred to as consolidation. This takes place mainly during sleep [17] and is believed to be effected by a combination of neuro-physiological mechanisms: Shortly after an initial episode of learning, cortical circuits undergo early tagging [18] and an immature engram is formed [19]. This is accompanied by a rapid growth of new dendritic spines [20, 21]. During sleep, the cortical engram is stabilized by replaying past neural activity [22–24] while task-irrelevant connections are pruned [21, 25, 26]. At the same time, surviving synaptic connections are collectively scaled down [27–29] in order to maintain firing rate homeostasis [30]. Notably, this regulation is multiplicative and, thus, preserves the relative differences between synapses [31].

Many of these aspects are neglected in standard attractor network models. Although phenomenological models have demonstrated that isolated aspects of consolidation, such as replay [32, 33], pruning [34], and homeostasis [35, 36], are beneficial for memory and learning, a principled account of the consolidation process within a common theoretical framework is lacking.

Here, we derive a normative synaptic plasticity model that reconciles the various biological mechanisms of consolidation with the notion of critical capacity in attractor networks. Our derivation is fundamentally based on a reformulation of the problem of critical capacity in two ways: First, instead of considering optimality to be a maximization of storage capacity [13–16], we define it as a maximization of memory robustness. Second, we assume that synapses are products of multiple sub-synaptic components which form the expression sites for synaptic plasticity [36–39]. The result is a self-supervised plasticity model that uses a combination of replay, homeostatic scaling and Hebbian plasticity to prune connections and shape the network to perform noise-tolerant memory recall. The model offers a simple explanation for a wide range of putative consolidation effects observed in synaptic, neural, and behavioral data.

## 2. Results

### 2.1. The circuit and synapse model

We model a local circuit of cortical pyramidal cells using a recurrent network of N excitatory binary neurons (Fig. 1a). At every discrete time step *t*, each neuron *i* = 1,…, *N* is characterized by an output state *s*_*i*_(*t*), which represents a brief period of elevated (*s*_*i*_ = 1) or suppressed firing (s_*i*_ = 0), similar to “up” and “down” states [40]. The elevated state (*s*_*i*_ = 1) occurs only if the neuron’s total input current *I*_*i*_(*t*) exceeds zero. This input current evolves in time according to

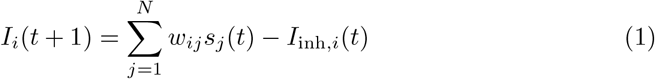

where the first term corresponds to the excitatory synaptic input from all neighboring neurons, with *w*_*ij*_ ≥ 0 denoting the connection strength from neuron *j* to *i*, while the second term summarizes the net effect of inhibitory inputs (see Methods 4.1).

**Fig. 1.**
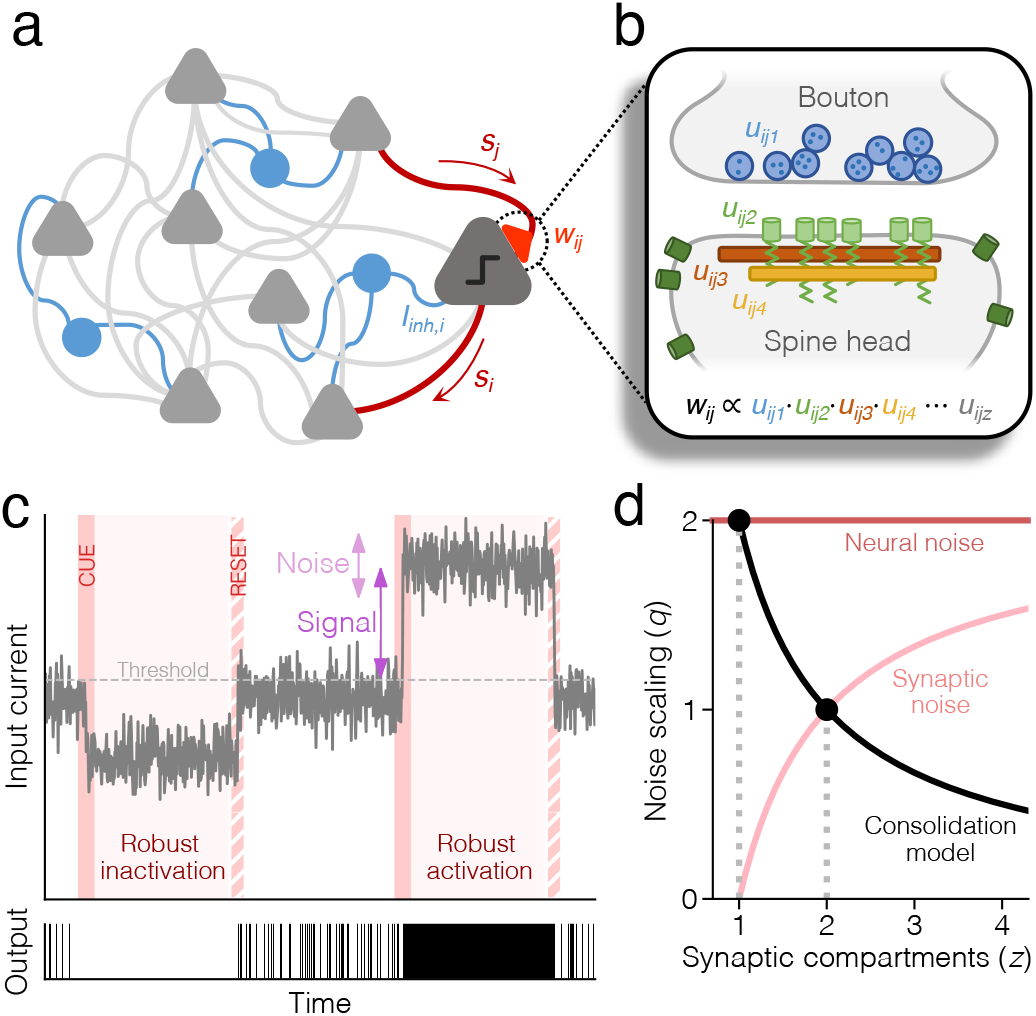
General model schematics. **(a)** Diagram of the circuit model. We consider a recurrent network of binary, excitatory neurons (gray) with non-negative connection weights, receiving a neuron-specific, scalar inhibitory input (blue). **(b)** Diagram of the synapse model. The total connection weight *w*_*ij*_ is a product of *z* factors *u*_*ij*1_,…, *u*_*ijz*_ that represent the efficacy of sub-synaptic components, e.g., release probability (blue), receptor density (green) and scaffolding protein content (brown, orange). **(c)** Illustration of input current dynamics during idleness (white background) and recall (pink background) in a single neuron. The SNR during recall of a pattern is determined by the deflection of the mean input current from threshold, relative to the fluctuations caused by noisy afferent neurons or synapses. **(d)** The noise scaling exponent *q* as a function of *z* for neural and synaptic noise. Consolidation with *z* components maximizes robustness with respect to noise of type *q* = 2*/z*, which is equivalent to neural noise when *z* = 1, and synaptic noise when *z* = 2.

In our mathematical analysis of the storage properties of the network, we focus on the connection strengths *w*_*ij*_. We begin by noting that the functional strength of a biological synapse (measured, for instance, as the amplitude of the excitatory postsynaptic potential, EPSP) is an aggregate quantity that is determined by the interaction of several protein complexes that combine to form the internal synaptic structure [41]. Following the induction of long-term plasticity, structural and chemical changes cascade throughout this molecular interaction network, causing the concentration and configuration of each component to be altered over the course of seconds to minutes [42]. This ultimately increases or decreases the synapse’s functional strength.

We model this internal synaptic structure by expressing each weight *w*_*ij*_ as the product of z internal, sub-synaptic components (factors) u_*ijk*_, where *k* = 1,…, *z*, so that

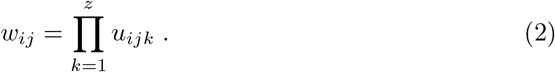

Each variable *u*_*ijk*_ can be seen as the relative concentration of a collection of one or more subcellular building-blocks that are necessary to form a functional connection, for instance, the average concentration of released neurotransmitters or the density of post-synaptic receptors and scaffold proteins (Fig. 1b; see Methods 4.2). Furthermore, consistent with the tagging-and-capture property [43, 44], we consider one of the synaptic components (*u*_*ij*1_) to be a flexible plasticity tag that is more volatile and sensitive to noise, while the remaining *z*−1 components are governed by more stable processes that are active only during consolidation.

### 2.2. Consolidation with homeostatic scaling, synaptic pruning, and replay

We define consolidation as the process of optimally storing a set of M memories, where each memory corresponds to a pattern of stationary network activity in which a specific group of neurons is active, while the rest is silent. The desired output of neuron *i* in pattern *µ* = 1,…, *M* is defined by 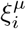, which is one with probability *f* ≤ 0.5 and zero otherwise. We parameterize the storage load using the ratio *α* = *M*/*N* (where *α*_*c*_ denotes the highest possible load).

Prior to consolidation, the network is assumed to have undergone an initial episode of learning that has imprinted all patterns as stable attractors, albeit with suboptimal robustness. At this stage, patterns can only be recalled if the network operates with very low levels of noise. The purpose of consolidation is now to tune all connections so as to maximize robustness and allow patterns to be successfully recalled under much noisier conditions.

We define robustness as the largest amount of noise that can be tolerated by the neural population before an error occurs during recall (Fig. 1c). This is determined by the signal-to-noise ratio (SNR) of the weakest pattern. We can optimize this by letting each neuron independently maximize a neuron-specific SNR where the signal is the amplitude of the input current deflection at the time the weakest pattern is recalled (see Methods 4.4). We write this as 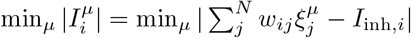.

The noise is determined by the magnitude of random fluctuations in the input current. This, however, varies depending on the noise source. Here, we expand on a previous analysis [45] and distinguish between two types of noise: neural noise and synaptic noise. Neural noise refers to perturbations of the network state (the s-variables) caused either by the encounter of distorted stimuli or by faulty neural output activity (i.e., firing below the threshold or failing to fire above the threshold). Synaptic noise, on the other hand, refers to perturbations in the connectivity, that, for example, are produced by spontaneous chemical reactions, conformational changes, or protein degradation and turnover [46, 47]. We model these perturbations as white noise added to the volatile u-component in each connection (see Methods 4.5).

Input fluctuations caused by neural noise scale as 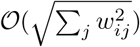 and are therefore dependent on synaptic weight but independent of synaptic structure. The magnitude of synaptic noise, however, depends both on synaptic weight and synaptic structure, by scaling as 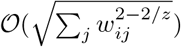 (see Methods 4.5). We can therefore write the SNR as

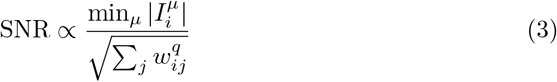

where we introduce a scaling exponent *q* which takes the value *q* = 2 for neural noise and *q* = 2 2/*z* for synaptic noise (Fig. 1d). The SNR can, in principle, be optimized (up to an arbitrary scaling factor) by any consolidation process that (a) maximizes the signal and (b) maintains a constant synaptic “mass” 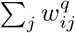. The latter property, however, necessitates a homeostatic weight regulation that is inhomogeneous across weights and, as such, directly at odds with the multiplicative homeostatic plasticity that has been observed experimentally [31] (see Suppl. Note S.1.3). We resolve this issue by optimizing the SNR in terms of each neuron’s sub-synaptic components *u*_*ijk*_, instead of directly treating the whole weight *w*_*ij*_ (see Methods 4.6). The result is the following three-step process (Fig. 2a):

**Fig. 2.**
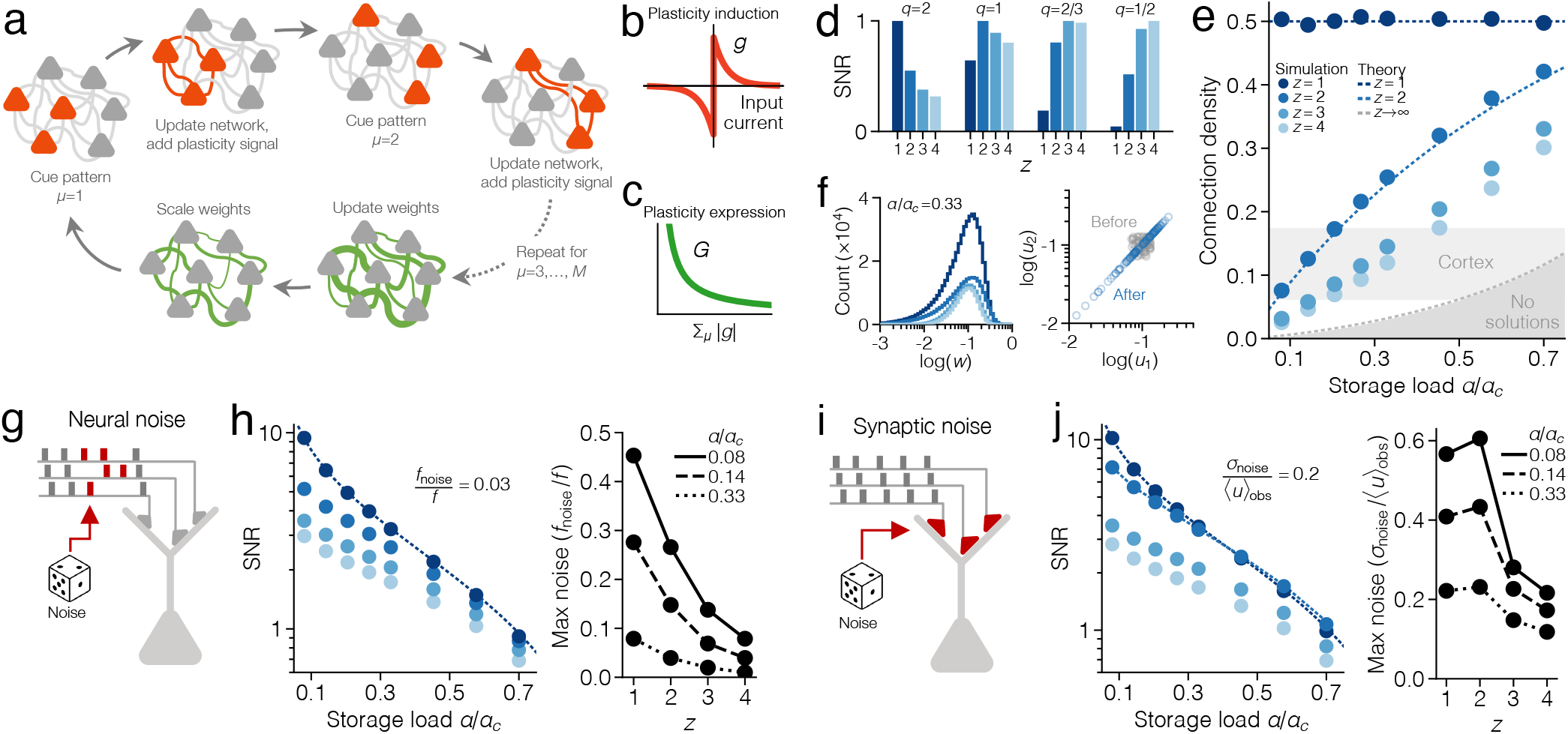
Simulated consolidation in networks with multi-factor synapses. **(a)** Diagram of one replay cycle of the consolidation model, implemented in discrete time. (**b**) The gating function *g*_*i*_. This determines the amplitude and sign of plasticity induction after replay of a single pattern. (**c**) The learning rate *G*_*i*_. This determines the amount of plasticity expression after a full replay cycle and depends on the accumulated signal ∑_*μ*_ |*g*_*i*_|. (**d**) SNR (mean over 10^3^ neurons) for different combinations of noise scaling *q* and components *z*, at *α*/ *α*_*c*_ = .08. Weights are normalized to 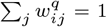, and the maximal SNR, for a given *q*, is scaled to one. **(e)** Connection density. Circles represent simulations (mean over 10_3_ neurons) while dashed lines represent theoretical solutions (Suppl. Note S.2). The light gray area marks the connection probability (mean ± SEM) among cortical pyramidal cells in a meta-analysis of 124 experimental datasets from mice, rats, cats, and ferrets [16] (Methods 4.12). **(f)** Left: distribution of weights (mean normalized to 0.1, colors as in e). Right: the second synaptic components (*u*_*ij*2_) plotted as a function of the first (*u*_*ij*1_) in a simulated neuron with *z* = 2, at *α*/ *α*_*c*_ = .33. **(g)** Illustration of neural noise. Each row of boxes represents binary input patterns at discrete time steps (gray = noise-free; red = distorted). **(h)** Left: SNR with respect to neural noise (*q* = 2; the noise level is parameterized by *f*_noise_; Methods 4.5). Right: highest level of tolerated neural noise in tests of pattern recall (Methods 4.7). **(i)** Illustration of synaptic noise, which directly perturbs synaptic strengths. **(j)** Left: SNR with respect to synaptic noise (*q* = 2−2/*z*; the noise level is parameterized by *“*_noise_). Right: highest level of tolerated synaptic noise in tests of pattern recall. All results in this figure are produced with f = 0.5, but there is no qualitative change with low-activity patterns (Suppl. Fig. S1).

- *Plasticity induction*: All patterns are replayed. For each pattern *µ*, the network receives a cue and is updated (Eq. 1) so that recall occurs. This triggers a plasticity signal 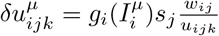 which is accumulated by the neuron. The *g*_*i*_-function is a neuron-specific, input-dependent plasticity gate that determines the sign and amplitude of induced plasticity (Fig. 2b; see Suppl. Note S.1.4).
- *Plasticity expression*: Once all patterns have been replayed, the accumulated plasticity signal is expressed by updating each component *u*_*ijk*_ with the increment 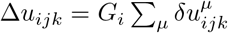, where *G*_*i*_ is a neuron-specific learning rate that is regulated so that the amount of expression is the same in each cycle (Fig. 2c; see Methods 4.6). Note that the fraction *w*_*ij*_/*u*_*ijk*_ implies that components that constitute a small part of their connection are more plastic, and vice versa.
- *Homeostatic scaling* : All *u*_*ijk*_ are scaled by a normalization factor, and the process starts over.

This consolidation model possesses a number of noteworthy mathematical properties: First, it is self-supervised, and requires no explicit error or target signal, as the target is provided by the response of the neurons themselves. Second, it maximizes the SNR with respect to noise with scaling exponent *q* = 2/z (Figs. 1d, 2d). Third, it is equivalent to *L*_2*/z*_-regularized optimization (see Methods 4.6), which means that a network with more sub-synaptic components prunes a larger fraction of its weights (Fig. 2e,f), despite the fact that homeostatic regulation always is multiplicative (regardless of z). Consequently, only networks with multi-factor synapses (z ≥ 2) reach a connection probability comparable to that measured in cortex (Fig. 2e). Fourth, the model forces components within a synapse to align with each other, so that *u*_*ij*1_ = *u*_*ij*2_ = … = *u*_*ijz*_ (Fig. 2f). All components therefore end up highly correlated with the total connection strength *w*_*ij*_, consistent with experimental findings [48, 49].

Networks with two-factor synapses (*z* = 2) are particularly important. While consolidation with *z* = 1 maximizes memory robustness with respect to *neural noise* (Fig. 2g,h), consolidation with *z* = 2 maximizes robustness with respect to *synaptic noise* (Fig. 2i,j). In practice, this means that two-factor consolidation generates networks that are highly pruned yet at least as robust to synaptic noise as the densest networks (Fig. 2j). From a neurophysiological perspective, these results are significant. When *z* = 2, we can describe the dynamics of the weights, close to convergence, with the differential equation

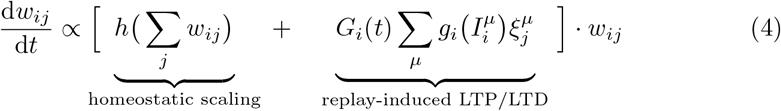

where *h*(*x*) is a general homeostatic function that is negative when x exceeds a baseline, and positive otherwise. All weight changes are now multiplicative, i.e., proportional to the momentary value of *w*_*ij*_. The homeostatic part, more specifically, performs a multiplicative *L*_1_-regularization that both prunes a large fraction of the connections and scales the remaining ones to maintain a constant average strength. This, by extension, keeps the average input current constant as well (assuming a stable level of output activity in the network). The formulation in Eq. 4 is directly compatible with, and generalizes, previously proposed models of homeostatic plasticity [35, 36] (Suppl. Note S.1.5).

Note that our consolidation model is entirely derived from normative assumptions. This is equally true for the synapse model in Eq. 2, which originates from a parameterization technique that implicitly biases an optimizer to find sparse solutions [50, 51]. Ablating either the sub-synaptic structure or the homeostatic scaling causes the model to fail (Suppl. Fig. S3).

### 2.3. Signs of consolidation in synaptic, neural, and behavioral data

In order to demonstrate how the consolidation algorithm can be incorporated into a single, self-supervised model of memory formation and stabilization, we simulate a network with two-factor synapses that optimally stores patterns across two phases of learning.

In the first phase, representing wakefulness, the network starts fully connected and sequentially encounters external stimulus patterns that are imprinted as attractors using few-shot learning (see Methods 4.9). This leaves the network densely connected and sensitive to noise (Fig. 3a). In the second phase, the network undergoes consolidation, rendering the connectivity sparse and robust (Fig. 3b; see Suppl. Fig. S4 for details). This process represents the cumulative effect of multiple sleep sessions taking place over an extended period of time.

**Fig. 3.**
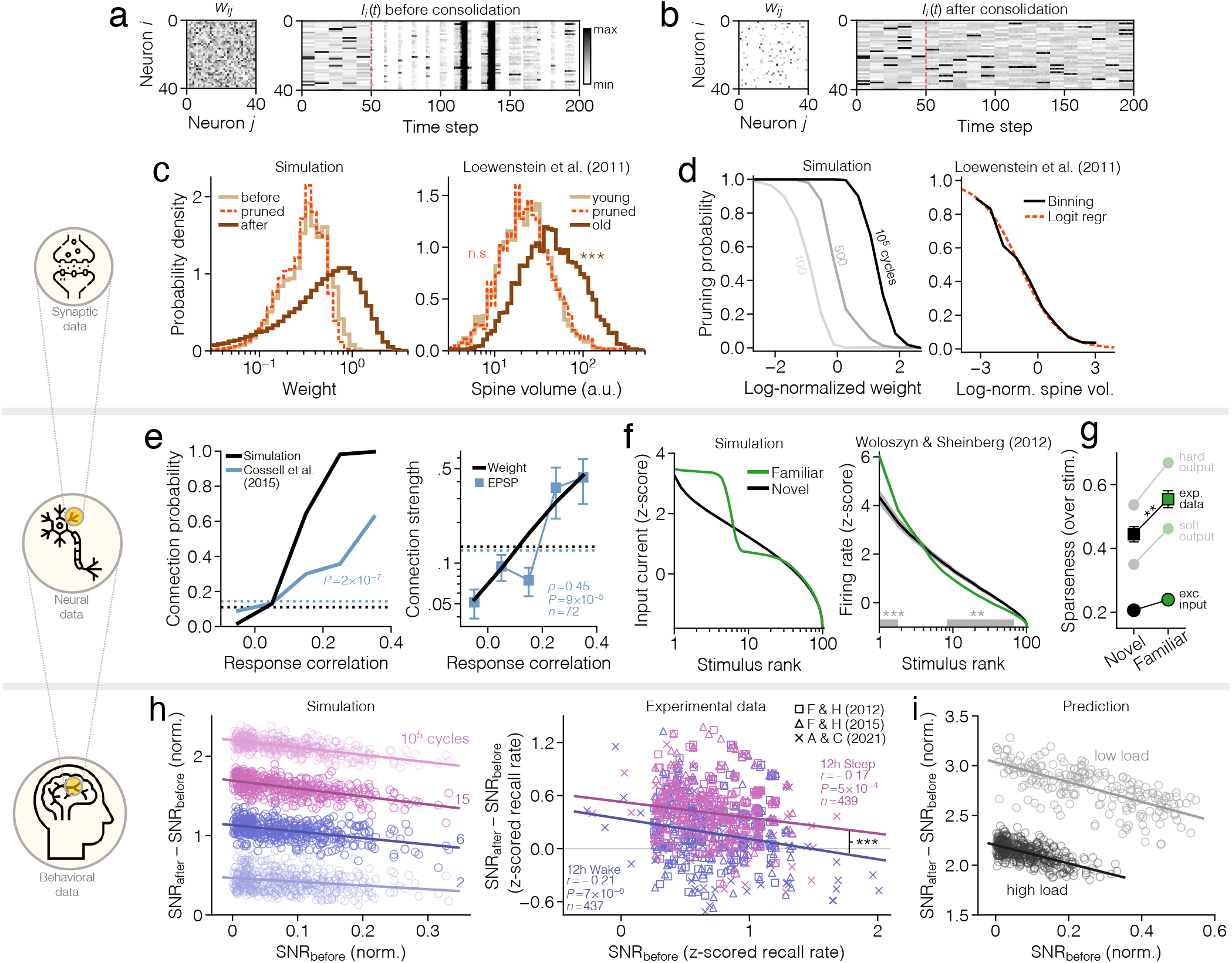
Signs of consolidation across three spatial scales. **(a)** Weight matrix (left) and input current (right) of 40 neurons during pattern recall, before consolidation (*f* = 0.05, *α*= 0.44). The network receives a cue every 10 steps and is then simulated for 10 steps. Synaptic noise starts after 50 steps (red line; *σ*_noise_/⟨u⟩_obs_ = 0.3). (b) Same as a, but after consolidation. (c) Distribution of weights (left) and dendritic spine sizes on pyramidal cells in rodent cortex [52] (right). (d) Pruning probability as a function of weight in simulated data (left) and as a function of spine size in experimental data (right). (e) Connection probability (left) and connection strength (right) as a function of binned response correlation among simulated neurons (black) and pyramidal cells in rodent visual cortex [53] (blue; error bars represent mean ± SEM). Dashed curves are grand averages. Connection strengths are normalized to have a maximum of one. (f) Tuning curves with respect to familiar and novel (previously unseen) stimuli, for simulated neurons (left; mean over 10_3_ neurons) and for pyramidal cells in macaque inferior temporal cortex [54] (right; mean ± SEM). (g) Tuning sparseness in simulations (circles; mean over 10_3_ neurons) and experimental data (squares; mean ± SEM). The hard and soft output is obtained by using sigmoidal activation functions with varying smoothness. (h) Left panel shows change in pattern SNR after simulated consolidation (circles are patterns) while right panel shows change in human memory trace SNR after sleep (pink markers) and after wake (blue markers) [55–57]. Behavioral data has been slightly jittered for clarity. (i) Change in pattern SNR after simulated consolidation with different loads. Stars indicate significance levels ∗∗*P* <0.01 and ∗∗∗*P* <0.001.

The simulation qualitatively reproduces a wide range of experimental observations linked to long-term plasticity (Fig. 3c-i; note, however, that simulated effects generally are more amplified, as we model a long stretch of biological time with a single bout of optimal consolidation). On the synaptic level, simulated wakefulness produces relatively small weight perturbations, while sleep entails more extensive rewiring. The distribution of pre-sleep weights therefore closely overlaps with the distribution of pruned weights (Fig. 3c, left), while surviving weights generally are stronger. We find analogous results in experimental data [52] (Fig. 3c, right). The distribution of dendritic spine volume for young spines (age 4*≤*d) is statistically indistinguishable from that of pruned spines, while old spines (age > 4 d) are significantly larger (Kolmogorov-Smirnov tests, *P*_pruned_ = 0.61, *P*_old_ = 8.6 × 10^*-*190^, *n*_young_ = 2268, *n*_pruned_ = 2300, *n*_old_ = 5011). This effect cannot be produced with single-factor synapses (Suppl. Fig. S5a).

An analysis of individual weight trajectories reveals that the probability of pruning decreases as a function of strength, meaning that connections that are potentiated prior to consolidation have higher chances of surviving (Fig. 3d, left). This trend is, again, present and highly significant in the experimental data [52] (logistic regression with two-tailed t-test, *P* = 1.5 10^*-*195^, *n* = 7311; Fig. 3d, right).

Next, we analyze how weights are configured depending on neural response similarities. Using the total excitatory input current ∑_*j*_ *w*_*ij*_s_*j*_ as an indicator of graded output activity, we find that neurons are more likely to stay connected after consolidation if their responses during recall are correlated (Fig. 3e, left). Similar synaptic selectivity is seen in experimental measurements of visual cortical neurons in mice during static image presentations [53] (two-sided Cochran-Armitage trend test, *P* = 1.7 ×10^−7^, *n* = 520). The average connection strength also increases with response correlation, both in simulated and experimental data (Spearman’s *ρ* = 0.45, *P* = 8.7×10^*-*5^, *n* = 72; Fig. 3e, right). Networks with single-factor synapses, however, fail to match experimental statistics (Suppl. Fig. S5b).

Another direct consequence of our consolidation model is an increased neural stimulus selectivity. Each neuron’s response to the stored patterns is enhanced by moving the input current further away from the threshold. This sharpens the tuning curve for familiar (consolidated) patterns relative to novel ones (Fig. 3f, left; see Methods 4.12). The same phenomenon can be observed in the activity of inferotemporal cortical pyramidal cells of Macaques, measured during the presentation of familiar and novel images [54] (Welch’s t-test, ^*∗∗*^*P* < 0.01, ^*∗∗∗*^*P* = 1.5*×*10^*-*5^, *n* = 73; Fig. 3f, right). The sharpness of the tuning curve is quantified by the *sparseness*, a metric that is near zero when all stimulus responses are similar, and near one when responses are selective to very few stimuli (see Methods 4.12). The sparseness increases significantly during stimulus familiarization (Welch’s t-test, *P* = 2.9*×* 10^*-*3^, *n* = 73; Fig. 3g).

On the behavioral level, sleep has been shown to enhance the ability to recall recently formed declarative memory [58], in a way that suggests larger improvements for items with weaker initial encoding [59]. We reproduce this effect by evaluating the change in SNR for each pattern over the course of simulated consolidation (Fig. 3h, left). Although a longer period of replay produces a stronger average encoding (curve shifts upwards), patterns that start off weak consistently benefit more than those starting strong (correlation is negative). This is a ceiling effect: as the SNR of each pattern is pushed to an upper limit, weak patterns inevitably exhibit a larger improvement than strong ones.

We further test the model by pooling and re-analyzing three large, published datasets on sleep-based consolidation of declarative memory [55–57]. In each study, humans memorize 40 word pairs and recall is tested before and after 12 h of wakefulness or sleep. We estimate the memory SNR in each subject as the z-scored recall rate, and then compute the change between the two test sessions. The result (Fig. 3h, right) confirms that gains in SNR are higher for subjects with weaker initial encoding, both after wakefulness (Pearson’s *r* = −0.21, *P* = 6.7*×* 10^−6^, *n* = 437) and sleep (*r* = 0.17, *P* = 4.6 *×*10^−4^, *n* = 439). There is no significant difference in the slopes (*t*-test, *P* = 0.49, *n* = 876), but sleep-gains are systematically higher across all initial performance levels (*t*-test, *P* = 3.9 ×10^−4^, *n* = 876). Our model predicts that a similar systematic shift in gains also should be observed when changing the word list length (Fig. 3i).

### 2.4. Implications for lifelong learning

To model the effects of consolidation over timescales of months and years, we start from the assumption that animals continually form new engrams throughout their lives, as a response to new and salient stimuli. This, in turn, increases memory load. We therefore represent cortical circuits at different stages in life using a network that has consolidated varying amounts of memory. We also use this model to represent cortical development under conditions of low or high environmental richness.

According to our model, a circuit that optimally stores a larger number of memories requires a higher density of connections (Fig. 4a, left). This is a direct consequence of maximizing SNR under sparseness constraints (see Fig. 2e, *z* ≥2). Importantly, it is consistent with the elevation in dendritic spine density that has been observed in animals raised in stimulus-enriched environments [60] (Student’s *t*-test, *P* = 5.7*×*10^−3^, *n*_low_ = 5, *n*_high_ = 6 for layer 5; *P* = 0.035, *n*_low_ = 4, *n*_high_ = 4 for layer 2/3; Fig. 4a, right). This experimental finding cannot be reproduced if we alter the consolidation model to maximize storage capacity instead of SNR, as has been suggested in past theoretical work [14–16, 62] (Fig. 4a, black; see Methods 4.11). The effect is also occluded when using single-factor synapses (Fig. 4a, gray).

**Fig. 4.**
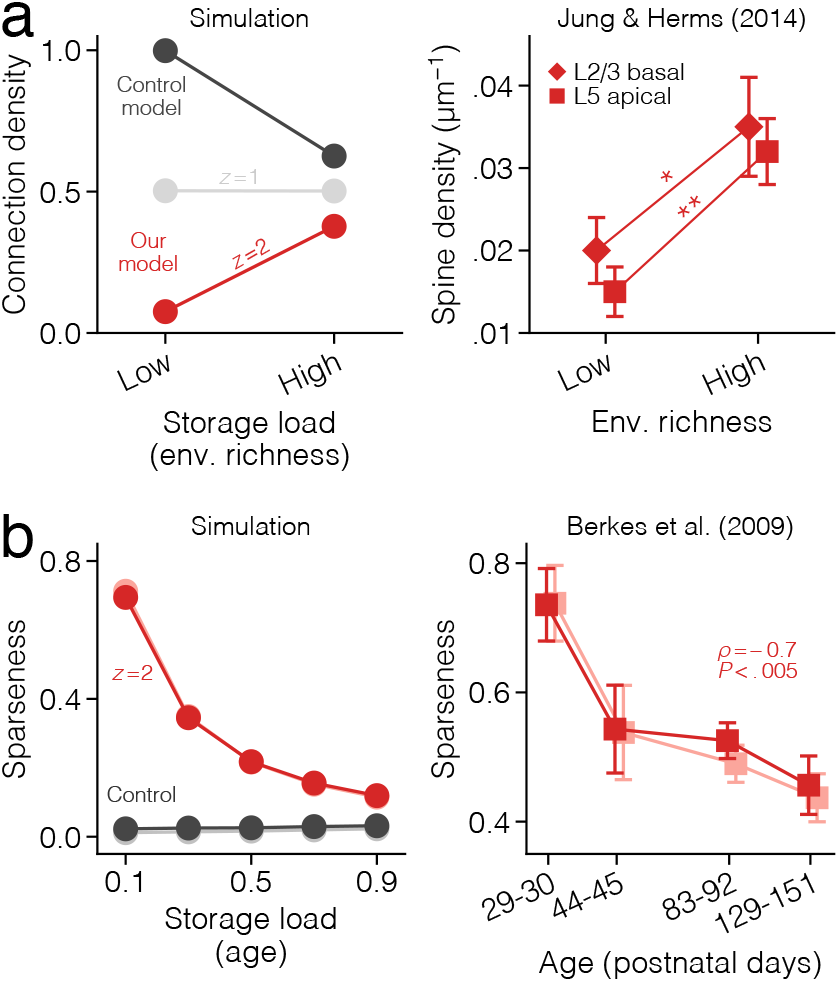
Signs of consolidation across development. **(a)** Left: Connection density as a function of storage load (indirect indicator of environmental richness) after consolidation with our model (z = 1, 2; same as Fig. 2) and with a control model that maximizes storage load instead of SNR (see Methods 4.11). Right: Density of stable dendritic spines (age > 3 weeks) in somatosensory cortex of rodents kept in environments of low and high stimulus richness since infancy [60]. Stars indicate significance levels ∗*P* < 0.05 and ∗∗*P* < 0.01. (b) Left: Sparseness across stimuli (red, black) and across neurons (pink, gray; see Methods 4.12) as a function of storage load (indirect indicator of age) after consolidating low-activity patterns (*f* = 0.05) with our model (*z* = 2) and the control model. Right: Sparseness across time (red) and across neurons (pink) for neurons in visual cortex of ferrets at different stages of development [61]. Circles represent mean over 10^3^ simulated neurons while squares represent experimental data (mean ± SEM).

Networks that optimally store more memories also exhibit flatter tuning profiles and, thus, decreased sparseness (Fig. 4b, left). This is a fundamental property of our consolidation algorithm, caused by the decrease in the maximum attainable SNR with load (see Fig. 2h,j). The effect is analogous to the decline in sparseness that has been measured in visual cortical neurons of ferrets at different stages of development, from eye-opening to adulthood [61] (Spearman’s *ρ* = −0.69, *P* = 2.9 *×* 10^−3^, *n* = 16 for sparseness across time; *ρ* = −0.67, *P* = 4.5*×*10^−3^, *n* = 16 for sparseness across neurons; Fig. 4b, right). This trend cannot be reproduced with a network that maximizes storage capacity instead of SNR (Fig. 4b, black).

### 2.5. Scaling of intrinsic synaptic noise

Our consolidation model crucially relies on the parameterization of each synaptic weight *w*_*ij*_ as a product of multiple components *u*_*ijk*_. Is it possible to detect signatures of such synaptic ultrastructure in available experimental data? To answer this, we first note that a key prediction of our model can be found in the synaptic noise scaling. When the volatile component *u*_*ij*1_ is subjected to random perturbations, the weight of the synapse, as a whole, fluctuates with an amplitude Δ*w*∝*w*^1−1*/z*^. For two-factor synapses, this reduces to

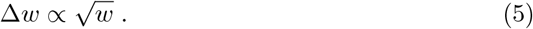

Stated more generally, our model predicts that synapses with more than one component display intrinsic noise that scales *sublinearly* with weight, both for potentiation and depression. It is only in the limit of infinitely many components (*z*→∞) that the noise magnitude becomes proportional to the weight. Conversely, only synapses with a single component produce intrinsic noise that is additive and uncorrelated with the weight.

To validate this prediction with an artificial synaptic dataset, we model the internal structure of a synapse as a stochastic dynamical system, and use this to simulate the evolution of 1000 independent synapses through time (see Methods 4.10).

The data is analyzed by plotting the absolute weight change | Δ*w*(*t*) |= |*w*(*t*+ Δ*t*) *−w*(*t*) | as a function of the initial weight *w*(*t*) and then applying a moving average to detect underlying trends in the scattered data (Fig. 5a). Consistent with our theory, the average noise amplitude, denoted ⟨| Δw|⟩, increases linearly in a log-log plot (Fig. 5a), both for depression (Δw < 0) and potentiation (Δw > 0). This indicates a power-law relation ⟨| Δw w^*x*^|⟩∝*w*^*x*^, where the exponent *x* is equivalent to the slope of the line in logarithmic space. We estimate this parameter using bootstrapped linear regression (see Methods 4.12), and find that it closely agrees with the theoretical prediction (Suppl. Fig. S6a).

**Fig. 5.**
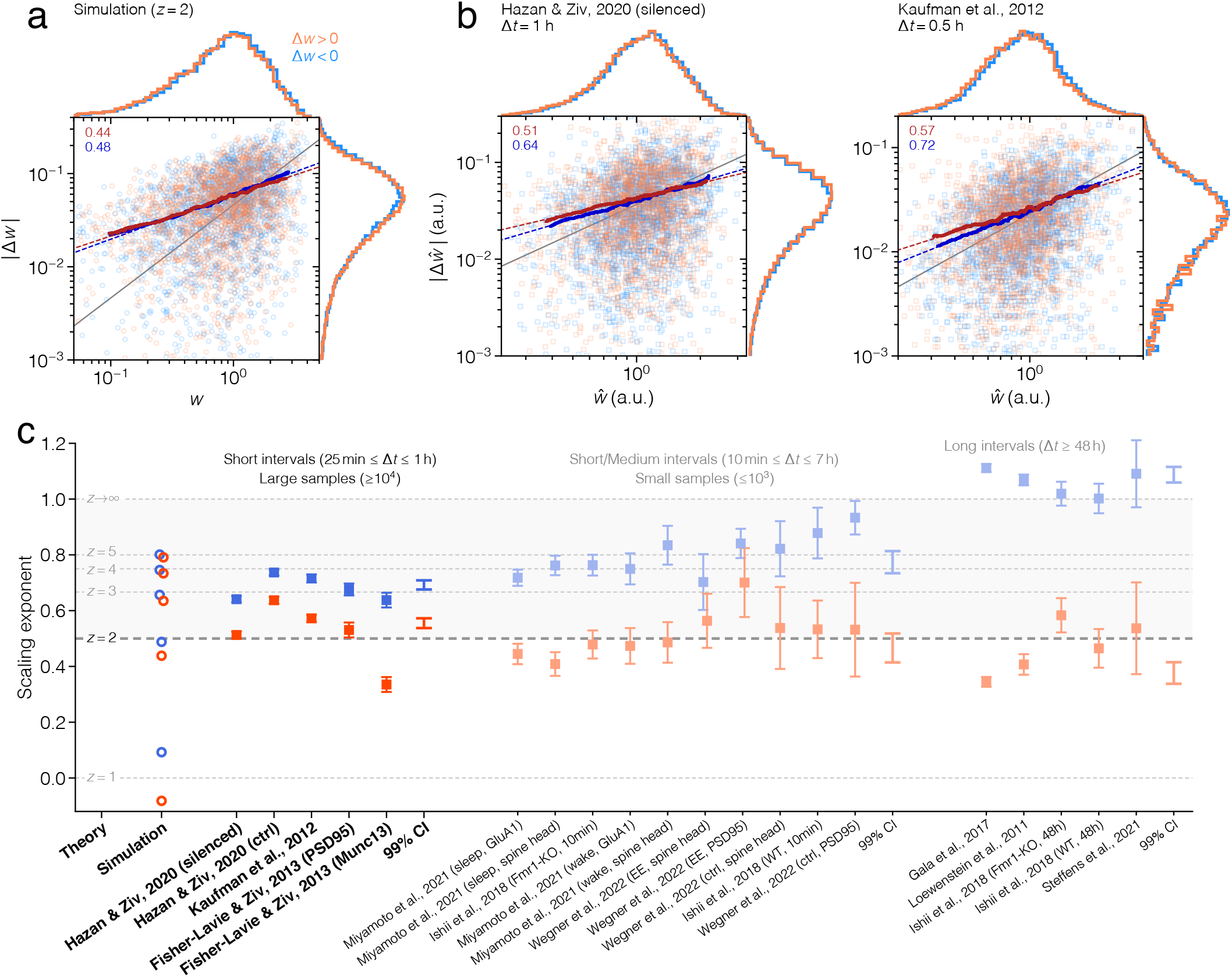
Scaling of synaptic fluctuations. **(a)** Absolute weight change as a function of initial weight in simulated data with z = 2, for potentiation (orange) and depression (blue); see also Suppl. Fig. S6a. Solid lines are the results of moving averages, and dashed lines are linear fits to the solid lines (slope value shown in upper left corner). The identity line (gray) has slope 1, and is included for comparison. (b) The same type of plot as in a, but for experimentally measured dendritic spine sizes in rodent cortical neurons [63, 64] (see also Suppl. Fig. S6b). (c) The scaling exponent of synaptic fluctuations in simulated (circles) and experimental data (squares; mean ± SE). This is the slope of the average fluctuation size in logarithmic space, obtained with bootstrapped linear regression. Labels on the abscissa contain a publication reference and a brief methodological descriptor; complete details are provided in Supplementary Tables S4, S5, and S6.

To test our prediction on experimental data, we compile 20 published synaptic datasets from 9 separate studies [29, 52, 63–69]. These publications span more than a decade of research and employ fluorescence microscopy and super-resolution nanoscopy in both cultured neurons and live animals, under various environmental conditions (see Suppl. Tables S4, S5, and S6 for details). Common to all studies, however, is that they measure an indirect indicator of synaptic strength (denoted 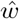) in a large population of synapses that have been individually tracked over extended periods of time (ranging from 24 h to almost 30 d).

We re-analyze each dataset according to the procedure described above. In the two largest datasets, shown as examples in Figure 5b, the average noise magnitude exhibits a clear linear dependence on the synaptic strength in logarithmic space, again indicating an underlying power-law like that found in simulations (similar results are reported in refs. 67, 70; see Suppl. Fig. S6b for more examples). The estimated noise scaling exponent for each dataset is presented in Figure 5c.

For large datasets with high sampling frequencies (i.e., short sampling intervals Δt ≤1 h; Fig. 5c, first group of data), synaptic fluctuations consistently have a sub-linear scaling, with an exponent of 0.56 *±* 0.02 for potentiation and 0.69 *±* 0.02 for depression (99% weighted confidence interval). These estimates are remarkably reliable and close to the range predicted by our synaptic noise model with *z* = 2 and 3. Note, however, that our model only describes *intrinsic* noise, which is best measured in conditions when activity-dependent synaptic plasticity is either negligible or entirely blocked. Theoretical predictions are therefore only approximately applicable to the experiments, which, in almost all cases, contain extrinsic synaptic noise. The data by Hazan & Ziv [64] is a notable exception, as this was acquired while glutamatergic transmission was pharmacologically blocked. In this case, the noise scaling almost exactly matches the theoretical lines for z = 2 and 3, as we obtain 0.51 *±* 0.01 for potentiation and 0.64 *±* 0.01 for depression (mean *±* SE, bootstrap of 100 samples; Fig. 5b, left).

In datasets with smaller sample sizes (Fig. 5c, second group) and longer sampling intervals (Fig. 5c, third group), the scaling exponent generally increases for depression and decreases for potentiation (Fig. 5c, second and third confidence intervals; see Suppl. Note S.4).

### 2.6. Signs of homeostatic scaling in synaptic noise

Our plasticity model does not only govern the trajectory of *individual* synapses, but it also shapes the distribution of synaptic *populations*. Recall that our model includes homeostatic scaling that, close to optimal storage, maintains a constant synaptic “mass” ∑w^2*/z*^. The implication is that the weight distribution, in the absence of activity-dependent plasticity, exhibits a constant 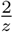 -th moment. To confirm this numerically, we return to the simulated synaptic data and estimate the stability of different moments of the weight distribution by calculating the coefficient of variation (CV) of the norm (∑w^*q*^)^1*/q*^ across time (Fig. 6b). Consistent with theory, we find that the weight norm that varies least over time (i.e., has lowest CV rank) roughly follows the relation q_min_ = 2/z (Fig. 6c).

**Fig. 6.**
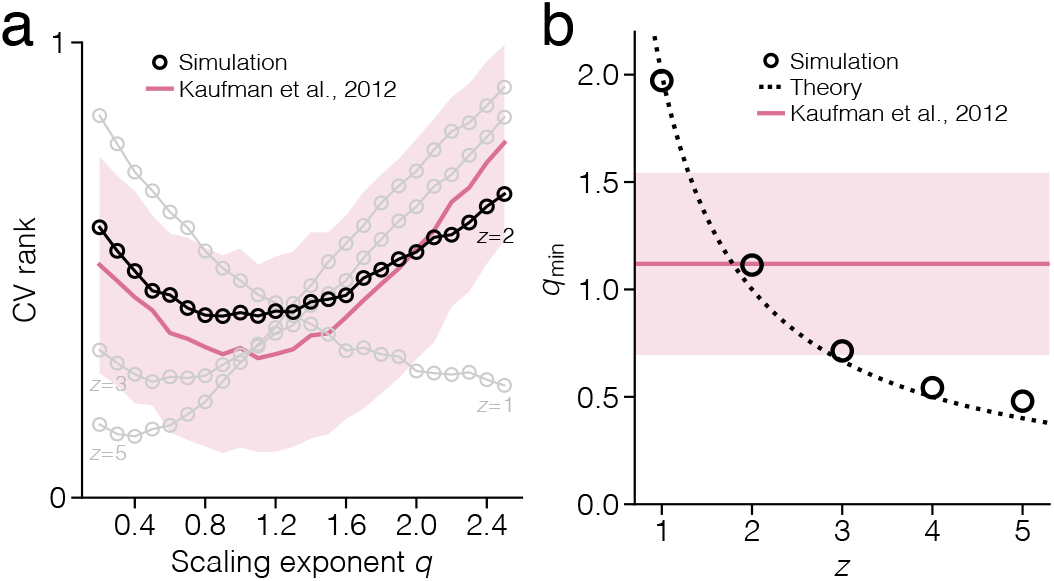
Signs of homeostatic scaling in synaptic noise. **(a)** The coefficient of variation (CV) of the norm (∑ w^q^)1/q, ranked from zero to one, as a function of *q* in simulated data (black, gray) and in dendritic spine sizes (red) measured on pyramidal cells from rodent cortex [63] (mean ± SE, bootstrap of 1000 samples). (b) The exponent qmin at which the CV in a is minimized.

We test this prediction using the experimental data reported by Kaufman et al. [63], which comprises 1087 dendritic spines, measured every 30 min over a total of 24 h. At each measurement, we calculate the norm of spine sizes, followed by the CV of the norm across time. The result, plotted as a function of q, displays a U-shaped curve that is best matched by the two-factor model (*R*^2^ = 0.20 for *z* = 2, compared to second best *R*^2^ = 0.14 for *z* = 3; Fig. 6b, pink curve). The smallest CV is obtained at *q*_min_ = 1.12 *±* 0.42, close to the theoretical prediction for *z* = 2 (Fig. 6c, pink line).

## 3 Discussion

We have derived a general mathematical model of synaptic consolidation based on the optimization of noise-robust recall of attractor memories in recurrent neural networks with factorized multi-component synapses. The contribution of our work is two-fold: First, it demonstrates that the various mechanisms underlying consolidation can be derived from first principles, within a single model of optimal memory storage. Second, by linking optimality to synaptic plasticity and the concept of critical capacity, it offers an explanation of how the structured connectivity of optimal attractor networks [14–16] might emerge in cortical circuits.

In the special case of two-factor synapses, our plasticity model takes a particularly simple form, in which all updates are multiplicative, both in terms of sub-synaptic factors u and the whole synaptic weight w. Despite this, a large fraction of all connections are pruned while the average strength of surviving synapses is homeostatically regulated. This resolves a contradiction in past synaptic plasticity studies: Sparse connectivity, like that measured in neocortex [6], has been difficult to reconcile with multiplicative homeostatic scaling [31], given that Hebbian plasticity with multiplicative constraints tends to produce dense solutions [71]. Sparse solutions typically require constraints that are either additive [16, 34, 71, 72] or that impose hard thresholds [34]. This, however, generally requires hyperparameter-tuning prior to learning (see Suppl. Note S.1.3). In our model, the introduction of sub-synaptic components reconciles the need for sparsification with multiplicative homeostatic plasticity.

Our results suggest that synaptic structural complexity serves a computational and metabolic purpose by implicitly biasing connectivity to be sparse, thereby lowering energy consumption and freeing unneeded synaptic resources for future learning. As such, our work is complementary to recent studies analyzing the effects of the synaptic ultrastructure on memory stability [73, 74], consolidation [43, 75], and energy consumption [76] (see also ref. 44).

We interpret our consolidation model as a general theory of sleep by situating it in the following scenario: During wakefulness, the network undergoes intense sensory-driven stimulation which imprints neural activity patterns as attractors. These are initially labile and, thus, represent immature engrams that are difficult to recall and are easily erased by spurious plasticity. During sleep, external inputs are silenced and patterns can be replayed. The process of consolidation now serves to tune connectivity in a way that enlarges all basins of attraction and pushes the network to critical capacity. This stabilizes the engrams and makes them resilient to structural and sensory perturbations.

Our model relies on a self-supervised replay mechanism that first reinstates memories sequentially, and thereafter modifies the synapses. This implies that memories must be recallable prior to sleep and that replay must be significantly faster than plasticity expression. Both requirements are supported by experimental observations [59, 77].

Our account of sleep-based consolidation offers an alternative to an earlier theory of sleep [78, 79], where replay is used to unlearn spurious attractors with anti-Hebbian plasticity in order to indirectly increase the robustness of desired memories. By contrast, our model is Hebbian and accomplishes the same goal by replaying only information that already is familiar, without having to identify spurious patterns.

The plasticity rule that forms the core of our consolidation model can be tested in synaptic, neural, and behavioral data. On the synaptic level, the model predicts that the internal structure of a synapse manifests itself as a sublinear scaling of intrinsic noise fluctuations. For two-factor synapses, this specifically means that noise scales as 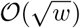. We emphasize, however, that an accurate analysis of synaptic noise requires a high sampling frequency and silencing of neural activity. Estimates of noise scaling are uninformative if the time between measurements is too long, as this only provides a temporal average that obscures the dynamics of instantaneous fluctuations.

On the neural level, our plasticity model requires a gating function that predicts that patterns linked to novel, immature, or otherwise weak memories induce higher levels of plasticity, compared to patterns representing highly familiar memories.

Finally, on the behavioral level, we predict that memory items that are weakly encoded prior to sleep generally display a larger improvement in SNR (and in the rate of recall) after sleep. While we partly confirm this with three large, published datasets, these cover only a part of the range of initial encoding. Moreover, our model predicts that the average recall performance should shift downwards when subjects are required to memorize more information, and vice versa.

We anticipate that our normative account of synaptic consolidation will contribute to a better understanding of long-term memory by inspiring neurobiologists to test the model with future experiments.

## 4. Methods

### 4.1. Circuit model

We model a local cortical circuit of pyramidal cells as a recurrent network of *N* binary neurons. At time t, the output state *s*_*i*_(*t*) of each neuron *i* = 1,…, *N* is given by

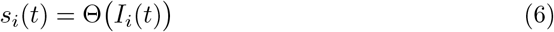

where Θ is the Heaviside function and *I*_*i*_ is the total input current, which is the sum of two non-negative current contributions, according to

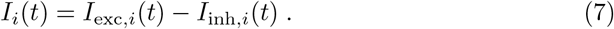

The first term is the excitatory input, which is determined by the recurrent connectivity and the previous state of the network, as in

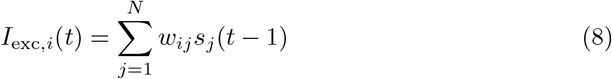

where *w*_*ij*_ *≥* 0 denotes the connection strength from neuron j to i. Self-connections are not allowed (i.e., *w*_*ii*_ = 0).

The second current term, *I*_inh,*i*_, is an inhibitory current which is neuron-specific and changes slowly, on a time-scale comparable to that of the excitatory weights (see plasticity rules below).

### 4.2. Synapse model

We consider each synapse to be comprised of *z* ∈ ℕ sub-synaptic components *U*_*ijk*_ (also referred to as factors), such that the strength of the connection as a whole can be written as the product

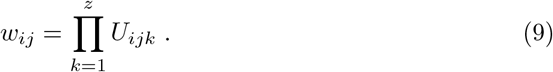

Each component can, for instance, be the area of a post-synaptic scaffold protein, a concentration of membrane receptors, a relative receptor efficacy, or a neurotransmitter release probability. It is therefore possible for each *U*_*ijk*_ to represent a separate type of physical quantity, with its own unit of measurement. In order to measure the strength of all components on a common scale, we rewrite each one as

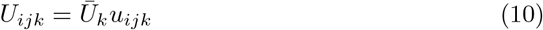

where *Ūk* is a constant that carries the unit and sets the measurement scale, whereas *u*_*ijk*_ is a unit-free measure that represents a relative strength on the same scale for all *k* (the measurement scale is implicitly defined by the constraint in Eq. 39). The weight can now be written as

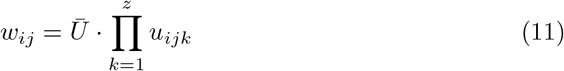

where the proportionality constant 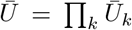 is the same across all weights and neurons. This constant only changes the length of all weight vectors, and can therefore be set to Ū = 1 without any loss of generality.

### 4.3. Memory patterns

Each memory pattern consists of a random binary vector 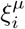, where *i* = 1,…, *N* is the neuron index, while *µ* = 1,…, *M* is the index of the pattern. Each element 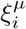 is independently assigned one with probability 0 < *f* < 0.5 and zero with probability 1−*f*. The parameter *f* is the average fraction of active neurons in each pattern, and is therefore referred to as the level of pattern activity.

We deviate slightly from this model when simulating wakefulness and sleep. In this case, each pattern contains *exactly fN* ones and (1−*f*)*N* zeros, to facilitate the few-shot learning procedure in wakefulness.

### 4.4. Memory robustness

The robustness of a single pattern *µ* with respect to neuron *i* is quantified with the signal-to-noise ratio of the input current at the moment of recall. We generally write this as

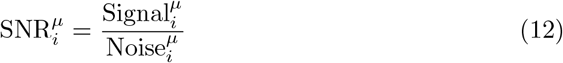

where both the signal and noise are pattern- and neuron-specific. As an approximation, we replace the noise with the strictly neuron-specific variant, by averaging across all patterns and obtaining

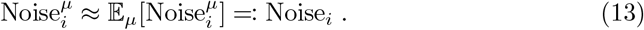

Expressions for this quantity can be found in the next section. The signal is calculated as the signed input current deflection during noise-free recall, that is

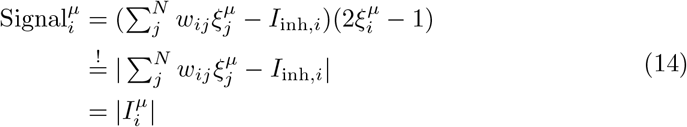

where the highlighted equality holds under the assumption that all pattern have been encoded error-free. This gives us the approximation

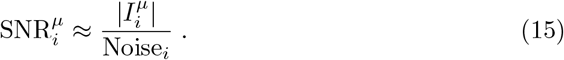

We now define the robustness of pattern *µ* as a whole as the smallest 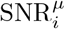 over all neurons, meaning

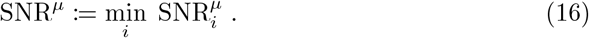

The robustness for multiple patterns, however, is ill-defined, as the optimization of SNR for one pattern can be incompatible with the storage of another. Therefore, in order to guarantee that no pattern is destabilized and forgotten, we define the total robustness of multiple patterns as the SNR of the weakest pattern, so that

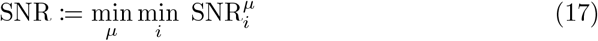

The order of the two minimizations can be switched. This enables us to maximize the total SNR by letting each neuron independently maximize its neuron-specific robustness

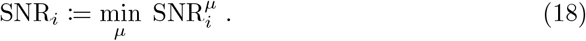

The optimal set of weights and inhibitions are defined as

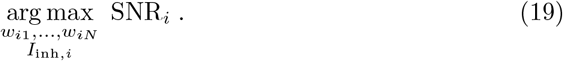

### 4.5. Noise scaling

One can distinguish between a total of three types of noise in the network: background noise, neural noise, and synaptic noise. We define the first two types in the same way as previous theoretical work [45], and then complement the analysis with the third type, which is new.

#### Background noise

Background noise refers to noise that is caused either by biochemical processes inherent to the neurons themselves, or by external inputs that are unrelated to the neural circuit we are observing. As such, we model background noise as a weight-independent, random current contribution *δ I*_*i*_ that is added to the total input current, according to

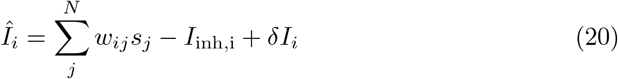

where 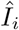 denotes a noisy, stochastic variant of the deterministic input current *I*_*i*_. Such noise can be made arbitrarily small in relation to the signal, irrespective of the tuning of individual weights, simply by scaling up the excitatory and inhibitory currents. We therefore omit background noise from further analysis.

#### Neural noise

Neural noise corresponds to noise that directly alters the output state of neurons. This is, for example, caused by distorted external stimuli or by transmission failures in afferent connections, that trigger firing when inputs are below threshold, or block firing when inputs are above threshold. We assume that distorted stimuli are generated by the same statistical process as the original patterns, and that they therefore retain the same average level of activity. To create a distorted instance of pattern *µ*, we flip the original pattern state 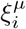 according to

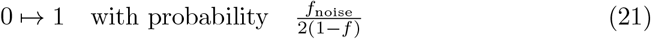

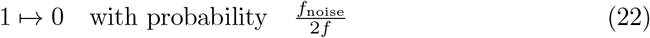

and obtain a new pattern 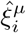, which, on average, contains *Nf*_noise_ errors, where *f*_noise_ is referred to as the noise level. This can be shown by calculating the expected error rate

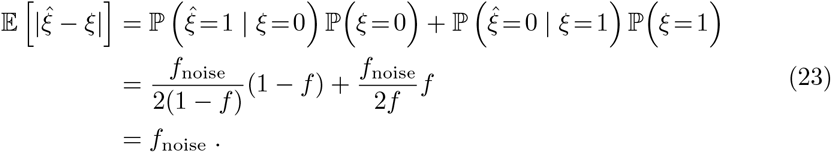

The activity level in the distorted pattern, however, remains unchanged, as shown by

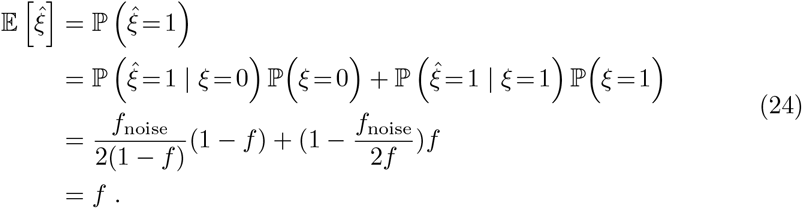

The distroted pattern 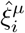 can be campactly described as a random variable

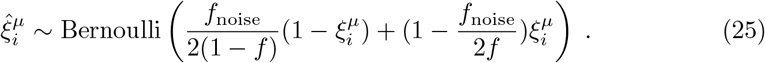

During pattern recall, we initialize each neuron *i* in the state 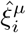, and update the network synchronously. Each neuron receives an input current that, across multiple trials, fluctuates with variance

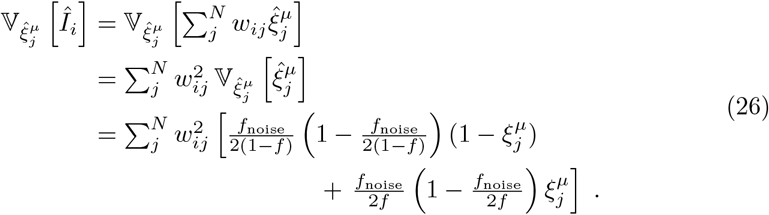

We average this quantity over all stored patterns and obtain

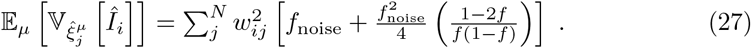

We finally estimate the noise fluctuation size as the averaged standard deviation

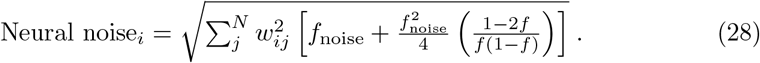

When evaluating the empirical robustness to neural noise, we report the results in terms of the relative noise level *f*_noise_/*f ≤* 2.

#### Synaptic noise

Synaptic noise represents intrinsic fluctuations in the most volatile constituents of the synaptic anatomy. We model this noise by adding a small, i.i.d. random perturbation *δu* to one of the sub-synaptic components in all observable (non-pruned) connections. The perturbation is drawn from a normal distribution 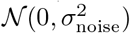. For simplicity, we assume that all sub-synaptic components are equal, so that *u*_*ij*1_ = … = *u*_*ijz*_ = *u*_*ij*_ (we later show that this assumption is justified in consolidated networks).

First, we note that a perturbation *δu* in one of the sub-synaptic components causes the whole weight to be perturbed with a magnitude *δw* given by

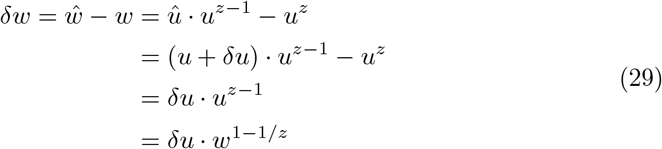

where we use the circumflex to, again, signify stochastically perturbed quantities. We test the impact of this noise on memory recall by first perturbing all connections in the network, then initializing each neuron *i* in a pattern 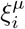, and finally updating the network synchronously. We separate the robustness analysis into two cases:

*z* = 1 After the first update, each neuron receives an input current that, across many trials, fluctuates with variance

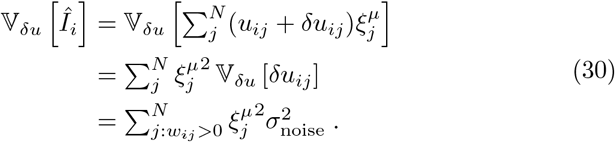

We average this quantity across all stored patterns and obtain

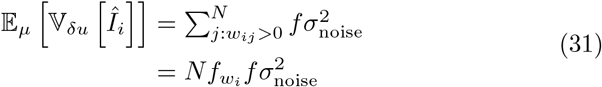

where 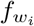 denotes the fraction of weights that impinge on neuron *i* and have not been pruned, that is

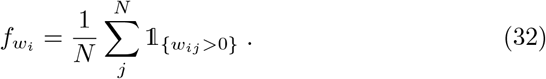

This yields the averaged standard deviation

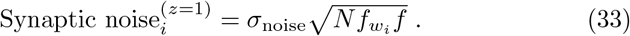

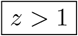 For multi-factor synapses, the variance of trial-to-trial input fluctuations is given by

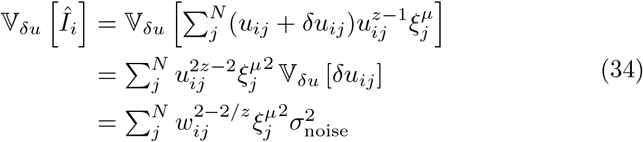

where we insert 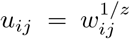 to produce the last expression. The average variance across all stored patterns is now

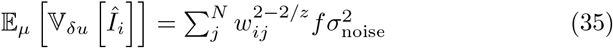

which yields the averaged standard deviation

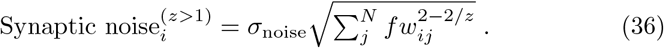

We compute the bias produced by synaptic noise as the difference between the average input current in the noisy and noise-free condition. This is zero for any *z*, as shown by

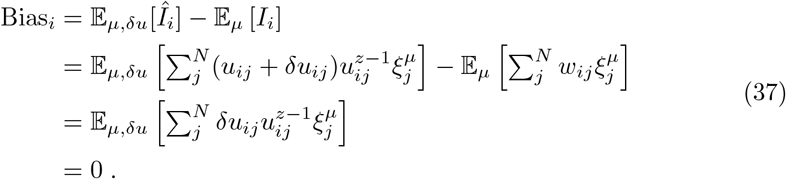

In order to compare the robustness to synaptic noise empirically across different network models, we always scale the noise level *σ*_noise_ relative to the mean of all observable synaptic components ⟨*u*_*i*_ ⟩obs in each neuron *i*, where

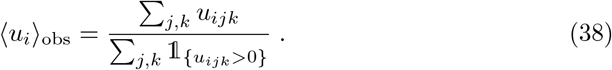

In practice, this is done by scaling all afferent connections so that ⟨*u*_*i*_⟩_obs_ = 0.1 prior to testing.

### 4.6. Consolidation algorithm

We define the process of consolidation as the maximization of the neuron-specific robustness SNR_*i*_ in each neuron i. We achieve this by maximizing the signal while keeping the noise fixed. The quantities *N, M, f, f*_noise_, and *σ*_noise_ are intrinsic to the circuit, and therefore considered constant. Crucially, we formulate the maximization in terms of sub-synaptic components u, to ensure that the resulting algorithm employs multiplicative homeostatic scaling. Thus, we formally define consolidation as the optimization

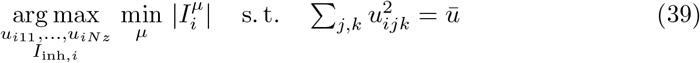

where Ū is an arbitrary constant. To make the problem more tractable, we denote the index of the weakest pattern as 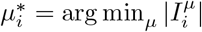 and rewrite the objective as

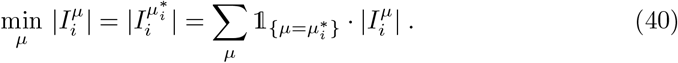

We solve this numerically using projected gradient descent. The derivative of the objective with respect to a weight component *u*_*ijk*_ is

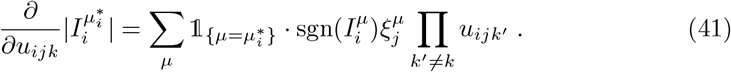

To avoid having to determine 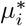 in practice, we replace the indicator function with its soft approximation, defined as

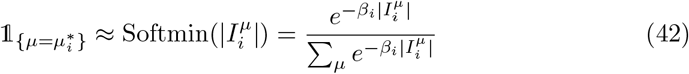

where *β*_*i*_ is a precision parameter (also referred to as an inverse temperature). This approximation becomes an exact equality in the limit *β→* ∞. Under the assumption that none of the sub-synaptic components are exactly zero, we further simplify the notation by writing

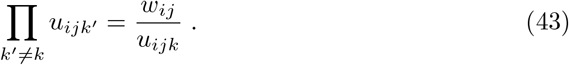

We now define a neuron-specific plasticity gating function *g*_*i*_ as

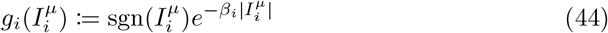

where we use a neuron-specific precision parameter *β*_*i*_, given by

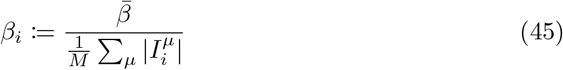

which adjusts the width of the gating function according to the average input current (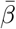 is a constant). We also use a neuron-specific learning rate

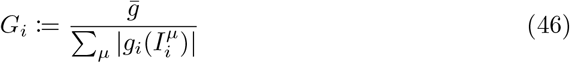

that is computed at the end of each replay cycle to ensure that the total sum of expressed plasticity stays roughly at a constant level (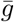 is a constant). We insert Eqs. 42–46 in Eq. 41 and obtain the synaptic update rule

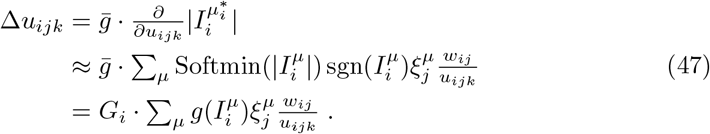

Analogously, the update for the inhibitory current becomes

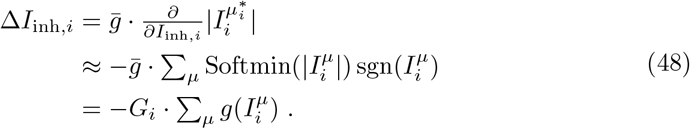

We summarize the discrete-time consolidation process in Algorithm 1. When *z* = 1, and with specific choices of the g-function, this algorithm reduces to the well-known gradient ascent, normalized gradient ascent [80], and batch perceptron algorithm [81] (Suppl. Note S.1.6).

At optimal weight configuration, all sub-synaptic components within the same weight adopt the same value, so that

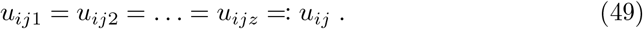

This is, in fact, a requirement that the solution *must* satisfy (see Supplementary Note S.1). The homeostatic constraint in Eq. 39 is now reduced to

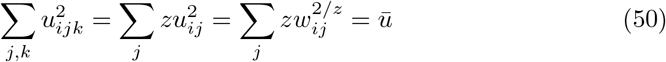

which means that the whole optimization problem is equivalent to an L_2*/z*_-regularized maximization, according to

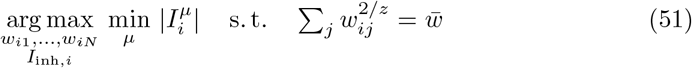

where 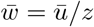.

#### Algorithm 1

Self-supervised consolidation in an attractor network

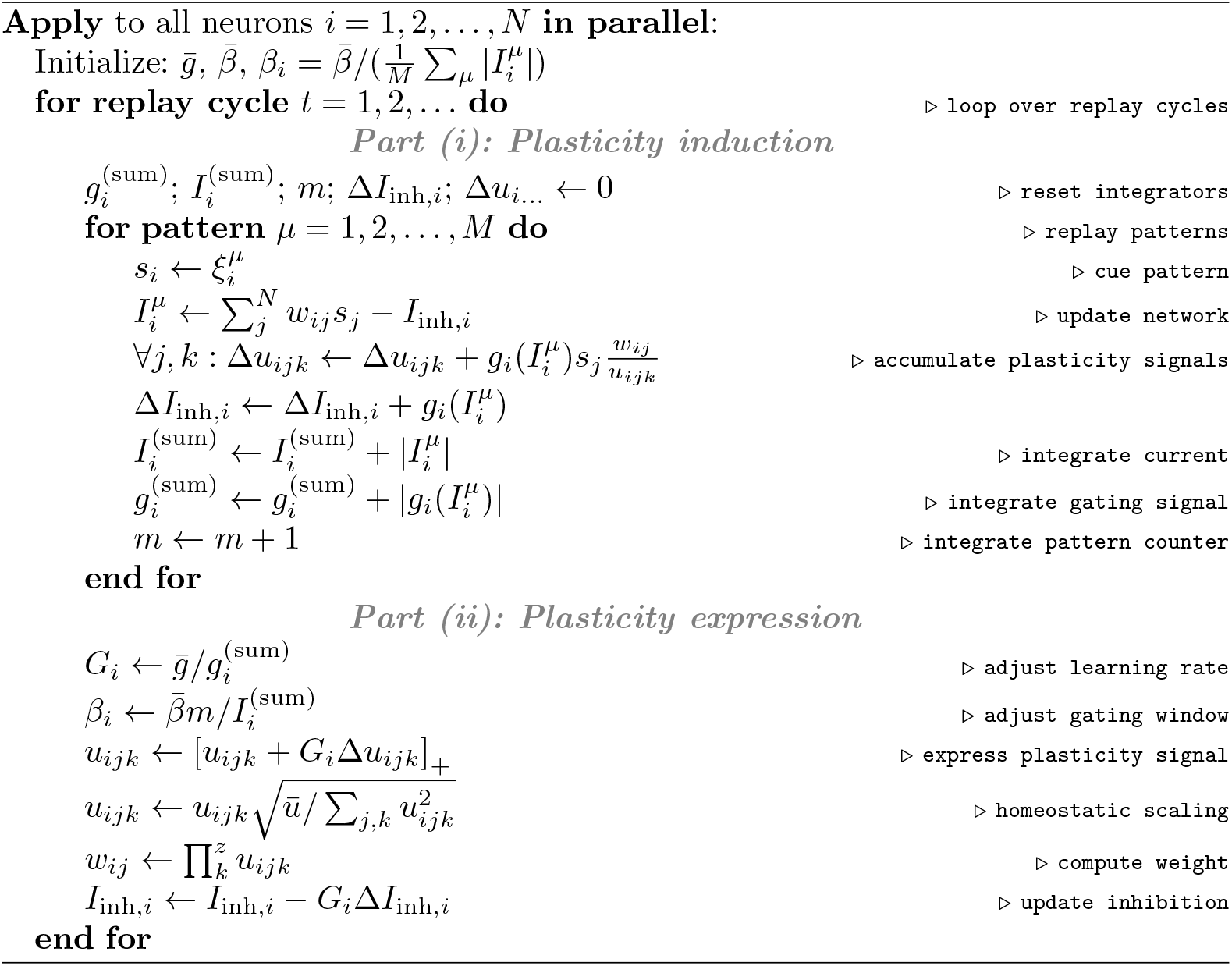

#### Continuous time

To study the dynamics of the weights in continuous time, we formulate the optimization in Eq. 39 as the penalized objective function

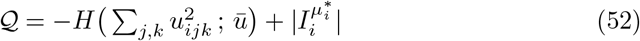

where 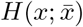 is a homeostatic penalty function that is zero only when 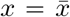 and increases monotonically everywhere else. The gradient is

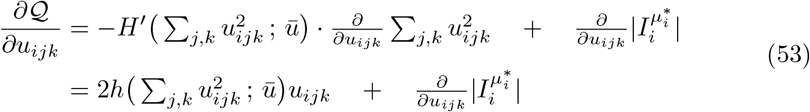

where *h* := −*H*^*′*^. We simplify the first term with the requirement in Eq. 50, and approximate the second term, as before, using Eqs. 42–46. Applying gradient ascent in the limit of infinitesimal learning rate gives us the gradient flow

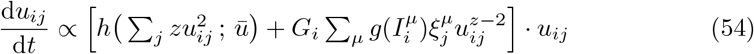

where a change of variables back to *w*_*ij*_ yields

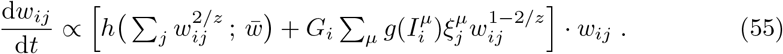

Note that both differential equations become multiplicative if, and only if, *z* = 2.

### 4.7. Numerical optimization and evaluation

#### Initialization

All sub-synaptic components are initialized by randomly sampling from the uniform distribution *𝒰* (0.7*u*_0_, 1.3*u*_0_), where *u*_0_ > 0 is an arbitrary constant. This ensures that the initial *u*-distribution is strictly positive and has a width that is 60% of the mean, regardless of scaling. Additional parameter values can be found in Supplementary Table S1.

In order to encode all patterns as attractors, prior to consolidation, the network is first trained using the batch perceptron algorithm [81] until all patterns can be recalled without error, where we define the recall error as the fraction of incorrect neurons after one synchronous state update. We compute this as

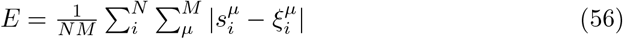

where

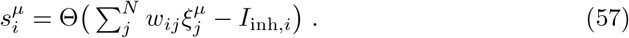

Once *E* = 0 is reached, the network is consolidated according to Algorithm 1.

#### Convergence

During the course of consolidation, we monitor the performance of the network using the average SNR_*i*_, average weight density 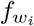, and error. The first two are calculated as

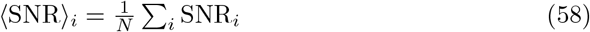

and

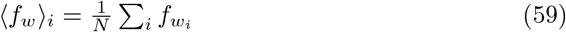

where the neuron-specific weight density 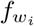 is estimated, in practice, using

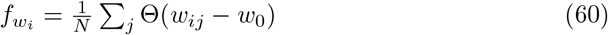

where *w*_0_ is the threshold at which a weight is considered pruned. In all simulations, we use *w*_0_ = 10^−10^.

We consider the consolidation to have converged once ⟨SNR⟩_*i*_ and ⟨*f*_*wi*_⟩change by less than 10^−4^ over 10^4^ replay cycles, while the error still is at *E* = 0.

#### Empirical robustness

After optimization, we empirically evaluate the robustness of the network by initializing it in each pattern *µ* together with either neural noise or synaptic noise. We then update the network 50 times and determine if the end state is close to the original pattern using the criterion that the error must satisfy E < 0.2f. We perform this test 20 times per pattern, with independent noise samples in each trial. We refer to the average fraction of patterns that can be recalled at each noise level as the recall ratio RR, and we define the empirical robustness as the noise level at which RR falls below 50% (Suppl. Fig. S1).

### 4.8. Theoretical solutions

The theoretical solutions in Figure 2 are adapted from previously published work, primarily references 11, 16, 82, 83. For more details, see Supplementary Note S.2.

### 4.9. Simulating wakefulness and sleep

We model fast, wakeful learning using a few-shot plasticity rule [84], gated by a novelty signal. In each replay cycle, every pattern is presented in random order to the network. This means that the network is initialized in a pattern *µ* and thereafter updated once. If the subsequent state of the network displays an activity level that differs from *f*, an additional inhibitory current 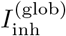 is triggered to regain the desired activity. This indicates that the pattern does not yet form an error-free attractor. The result is registered by the novelty signal 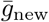 according to

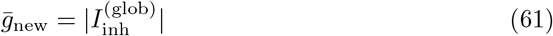

and the network is initialized once again in pattern *µ* and updated according to

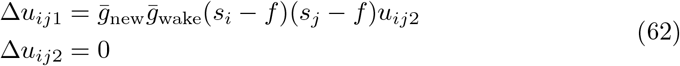

without any homeostatic scaling. Inhibition is adjusted to balance excitatory input according to

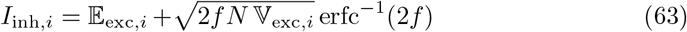

where 𝔼_exc_ and 𝕍_exc_ is shorthand for the mean and variance of the excitatory input current across pattern presentations, calculated as

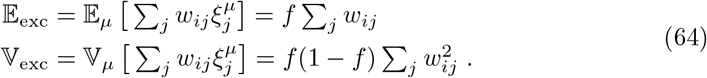

Wakeful learning is repeated until none of the patterns trigger the novelty signal. At this point, sleep commences, and both *u*_*ij*1_ and *u*_*ij*2_ are allowed to change. Consolidation is now modeled using Algorithm 1. Parameter values can be found in Supplementary Table S2.

### 4.10. Simulating synaptic intrinsic noise

To simulate synaptic noise, we assume that one sub-synaptic component is volatile and changes with a fast time constant (fixed to 1), while all remaining components are more stable and characterized by the time constant τ *»* 1. Each weight is therefore parameterized as

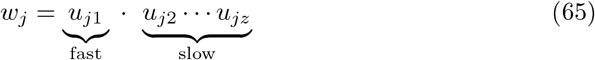

where *j* = 1,…, *N*. The fast and slow components are governed by the stochastic dynamical system

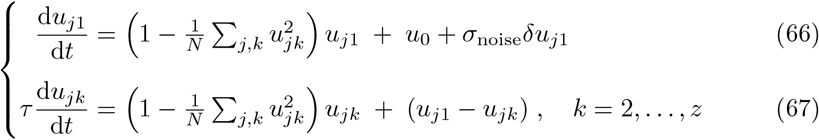

where δ*u*_*j*1_ *∼ 𝒩* (0, 1), *u*_0_ is a bias, and *σ*_noise_ scales the amplitude of the noise fluctuations. All components are initialized at *u*_*jk*_ = 1 and simulated with step size dt = 0.005 for a total time of *T*_sim_ = 10^3^, with a sampling time of *T*_sample_ = 1. The analysis in Figures 5 and 6 is performed using the last 144 samples (which corresponds to approximately 24 h if the time unit is assumed to be in the order of 10 min). Additional parameter values can be found in Supplementary Table S3.

### 4.11. Control model

In Figure 4, we compare our model with a control model that has been used in past publications to train attractor networks to achieve optimal storage [13, 15, 16, 62]. The latter approach is based on the assumption that cortical circuits store patterns with an SNR that is inherently fixed by the plasticity model. Early in development, the storage of new stimuli increases the load of the circuit (as long as *α* < *α* _*c*_), until critical capacity is reached (*α* = *α* _*c*_), at which point the circuit enters a steady state where additional storage of new patterns is counterbalanced by forgetting old ones [14]. Stated mathematically, this type of consolidation maximizes the storage of the network at a fixed SNR, by solving

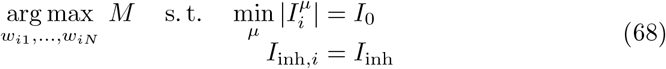

where *I*_0_, *I*_inh_ > 0 are constants, and no further reparameterization of the weights is used. The first condition ensures that the signal is fixed, while the second condition imposes a constant inhibition. In ref. 13, it is shown that the solution to Eq. 68 satisfies

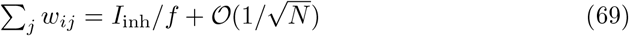

which means that the sum of the weights is constant, up a to correction term that vanishes as *N* →∞. If both the signal and the summed weights are constant, then the SNR with respect to *q* = 1 is also constant, which we write as

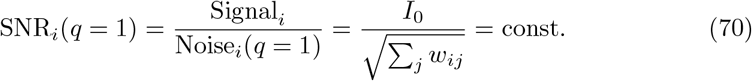

We emphasize that this differs from our consolidation model, where we instead maximize the SNR for a fixed storage load *M* (see Eq. 39).

Borrowing the notation in reference 13, Eq. 70 is equivalent to a fixed robustness parameter

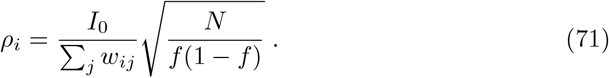

We train the control model using a variant of the perceptron algorithm, whereby we present each pattern *µ* to the network and compute 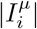 for every neuron *i*. The weakest pattern in each cycle is tagged with index 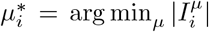 and used to calculate the robustness *ρ*_*i*_. The neuron’s weights are now updated according to

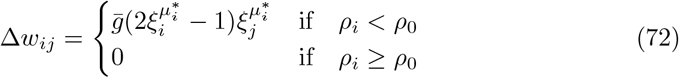

where *ρ*_0_ is the robustness threshold. The process is repeated until at least 99% of all neurons satisfy *ρ*_*i*_ *ρ*_0_, at which point the optimization stops.

### 4.12. Data analysis

#### Cortical connectivity

The experimental data on connection probability among cortical excitatory cells is part of a publicly available compilation of 124 datasets that were included in a meta-analysis published by Zhang et al. [16]. We assign each dataset a weight *β* according to the number of evaluated potential connections *n*_conn_, so that

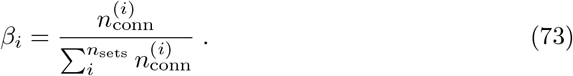

The weighted mean (wM) and weighted standard error (wSE) of the connection probability P_conn_ is estimated using

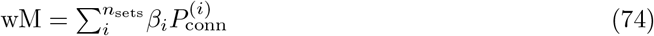

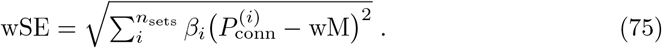

#### Synaptic pruning

To analyze the properties of synaptic pruning, we utilize the dataset published by Loewenstein et al. [52]. This consists of dendritic spine volume measurements conducted across six sessions, separated by a sampling interval of Δ*t* = 4 d (see Table S6 for details). We separate spines into three categories: (i) Spines that are first observed sometime between sessions 2 and 6 are defined as “young”. Spines observed in session 1 have an unknown age, and are therefore left out. (ii) Spines that disappear at any time between sessions 1 and 6 are defined as “pruned”. (iii) Spines that can be seen in at least two consecutive sessions are defined as “old”.

To estimate the pruning fraction, we first log-normalize the data by calculating the z-score in logarithmic space, according to

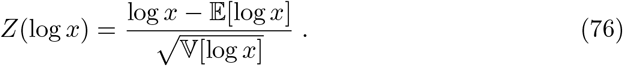

We then bin all spine volumes in sessions 1 to 5, and compute the ratio between the number of pruned spines and the total number of spines in each bin. Spines in session 6 are omitted, as it is unknown how many of these that are pruned.

We calculate the simulated pruning fraction in the same way, by comparing connection weights that are pruned during sleep to all connection weights before sleep.

#### Connection selectivity

In order to evaluate how network connectivity depends on neural response properties, we use the excitatory input current during pattern recall as a proxy for graded neural activity, and denote this 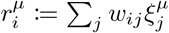. We use this to calculate the neural response correlation between two neurons *i* and *j* as

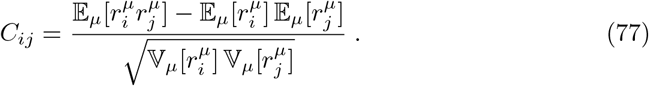

To estimate the connectivity and connection strength as a function of response correlation, we bin all neuron pairs according to *C*_*ij*_ and thereafter compute the connection probability and average weight in each bin.

We compare our simulations with the experimental data published by Cossell et al. [53]. This study reports the connectivity among pyramidal cells in layer 2/3 of mouse visual cortex, together with their neural activity and pair-wise correlations during presentations of natural static images. The authors estimate the connectivity and synaptic strength (in terms of excitatory post-synaptic potentials, EPSPs) as a function of pairwise correlations by binning neuron pairs as described above. In order to compare artificial weights with biological synapses, we normalize all weights and all EPSPs with the largest value in each dataset.

#### Stimulus tuning

We compute the neural response to a familiar (i.e., consolidated) pattern *µ* using 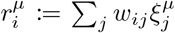 Analogously, the response to a novel pattern is computed as 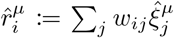, where 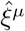 denotes a previously unseen pattern that is created by randomly shuffling all entries in pattern *ξ*^*µ*^. In order to produce the tuning curve, we first z-score the response distribution of each neuron relative to its familiar responses, according to

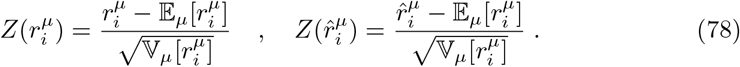

and we then sort all Z-scored responses and plot them as a function of their rank, ranging from 1 (highest) to 100 (lowest).

The sharpness, or selectivity, of the tuning is quantified with the *sparseness* [85, 86], which is defined in general terms as

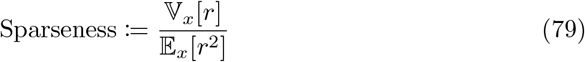

where *r* is a general neural output activity (e.g., firing rate). This is computed either across stimuli (*x* = *µ*) or across neurons (*x* = *i*); the former variant is typically called *lifetime sparseness*, and describes the selectivity of single neurons, whereas the latter is called *population sparseness*, and describes the response to a single stimulus in the entire population (see Suppl. Note S.3 for more details).

We compare the simulated results with the data published by Woloszyn and Sheinberg [54]. This consists of firing rates measured in putative excitatory neurons in inferior temporal cortex of macaque monkeys during presentation of familiar and novel images of objects. The experimental firing rates are processed in the same way as the modeled neural responses.

#### Associative memory tests in humans

In order to estimate how the strength of memory encoding changes across wakefulness and sleep, we utilize the behavioral data reported by Fenn and Hambrick [55, 56], and Ashton and Cairney [57]. All three studies involve human subjects tasked with memorizing 40 semantically related word pairs, where recall performance is tested before and after a delay of roughly 12 h of wakefulness or sleep. We model the recall process according to signal detection theory [87], by assuming that the trace of a memory is encoded in each subject according to a subject-specific strength that is perturbed by noise at encoding time. All traces within a subject are therefore assumed to be approximately normally distributed after the initial training session. During testing, only memories whose trace exceeds a subject-specific threshold can be correctly recalled. We define the recall ratio as

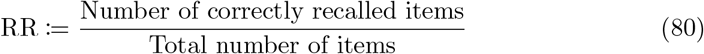

and use this to estimate the average memory SNR in a subject as the distance from the average trace strength to the threshold; this is given by the z-scored recall ratio

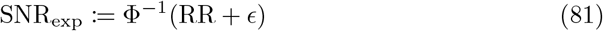

where Ф is the normal cumulative distribution function and ϵ = (1 −2RR) 10^*−*16^ is a small corrective term added to avoid divergence. We calculate the change in SNR over the course of the delay period as

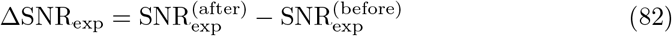

and pool the three datasets. Data points that are further than four standard deviations from the mean are considered outliers and are removed. The data is then fit with the linear model

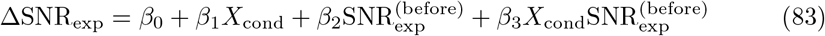

where the experimental condition is coded by the categorical variable

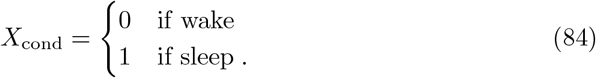

We determine if the intercept and slope differ significantly between wake and sleep by conducting a one-sample *t*-test of (3_1_ and (3_3_ relative to zero.

#### Environmental enrichment

In order to analyze the effects of environmental enrichment on cortical connectivity, we reference the study by Jung and Herms [60]. This dataset contains measurements of dendritic spine density in the somatosensory cortex of mice that are kept in either stimulus-enriched or stimulus-impoverished environments from birth to adulthood. We reproduce the density of spines that are classified as “persistent”. These are older than 3 weeks and are therefore part of connections that, presumably, have undergone maturation and stabilization.

#### Sparseness throughout development

To observe how neural activation sparseness changes over long periods of time, we use the experimental data reported by Berkes et al. [61]. This consists of spike-time measurements in the visual cortex of awake ferrets that are shown a movie clip at different stages in development, ranging from the period of eye-opening to adulthood. We calculate firing rates by binning the spike data in 10 ms bins. The sparseness is then obtained using Eq. 79.

#### Synaptic noise scaling

To study the scaling of synaptic noise, we use 20 different datasets of synaptic measurements, acquired in 9 previously published studies [29, 52, 63–69]. In general, each datapoint consists of a measurement of a synaptic strength proxy, denoted 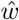, and the observed change 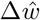 following a sampling interval Δ*t*. We first separate the data into potentiation 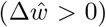 and depression 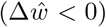 and then calculate the average absolute change 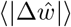 as a function of initial strength by filtering all datapoints in 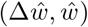,-space with a moving average, using window size n/20, where *n* is the sample size.

We obtain an estimate of the scaling exponent as the slope of 6.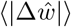 in logarithmic space, using linear regression. The mean (M) and standard error (SE) of the exponent is estimated by repeating the averaging and line-fitting with bootstrapping. All datasets are bootstrapped 1000 times, except the datasets in references 63–65, which are bootstrapped 100 times due to their exceptionally large sample size.

To summarize the estimates across datasets, we assign each estimate *i* a weight *β* according to its inverse variance (squared standard error), as in

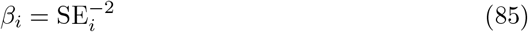

and we use this to calculate the weighted mean (wM) and weighted standard error (wSE) according to

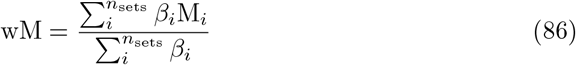

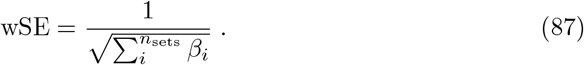

The 99% confidence interval is finally estimated as [wM *±* 2.58 *·* wSE].

#### The CV of synapse norms

We use the artificial synaptic data that is obtained by simulating the dynamical system in Eq. 66, and we analyze only weights that survive until the end of the simulation (i.e., *w*_*j*_(*T*_sim_) > 0). At each sampling time t, we calculate the q-norm of the weights with

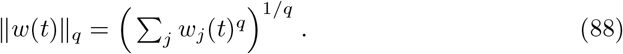

We then compute the CV of the q-norm across samples, according to

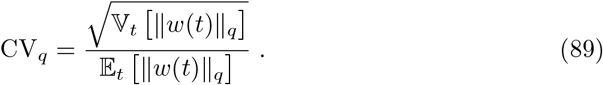

After repeating this process for a range of q-values, we compute the CV-rank by rescaling all CV_*q*_-values to lie in the range [0, 1] (from smallest to largest) and we obtain the norm with smallest CV as

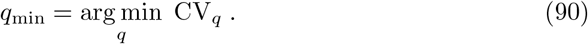

We estimate the mean and standard error of the CV-rank and q_min_ by bootstrapping this procedure 1000 times. In each run, we generate the bootstrapped data by separately re-sampling weights at each time *t*.

We compare simulated results with experimental data by utilizing the dendritic spine measurements reported by Kaufman et al. [63]. The experimental CV-rank and *q*_min_ are computed in exactly the same way as for the artificial data.

## Supplementary information

All Supplementary Notes, Figures, and Tables can be found in the Supplementary Material. The simulation code can be found at github.com/geoiat/2f-syn-con.

## Acknowledgments

The authors would like to thank Prof. Haruo Kasai, Prof. Noam Ziv, Prof. David Sheinberg, Prof. József Fiser, Prof. Maria Florencia Iacaruso, Prof. Kimberly Fenn, Prof. Armen Stepanyants, and Dr. Rohan Gala for sharing their experimental data.

This study was supported by funding from the Swiss government’s ETH Board of the Swiss Federal Institutes of Technology to the Blue Brain Project, a research center of the École Polytechnique Fédérale de Lausanne (EPFL).

## Author contributions

GI created the model, performed the simulations, and analyzed the data. JB assisted in the data analysis and theoretical derivations. GI, JB, and WG wrote the article.

## Supplementary Material

### S.1. Analysis of the consolidation algorithm

In this section, we explain the mathematical foundation of our consolidation algorithm and clarify its relation to other learning algorithms in the literature. As in the main text, we consider a recurrent neural network of *N* binary neurons with inhibition *I*_inh,*i*_ and connection weights *w*_*ij*_ 0, where *i, j* = 1,…, *N*. For the sake of brevity, we introduce vector notation and represent all input weights to neuron *i* with the column vector ***w***_*i*_ = (*w*_*i*1_,…, *w*_*iN*_) ^⊤^ and each pattern *µ* as 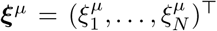 For added simplicity, we omit subscript *i*. The definition of memory robustness in Eq. 3 can now be expressed as

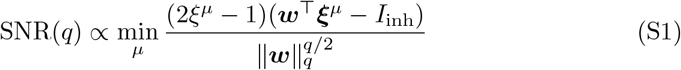

where the exponent *q* > 0 is chosen depending on the type of noise that is considered. Likewise, the aim of consolidation, as stated in the main text, can be written as the neuron-specific optimization

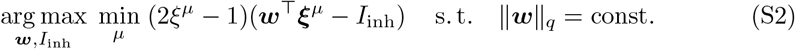

This maximizes SNR(*q*) subject to a homeostatic constraint placed on ║ ***w*** ║_*q*_. Note, however, that without such a weight constraint, the SNR has no upper limit (for *q* < 2) and can be scaled up indefinitely, at a rate *c*^1−*q/*2^, simply by scaling the weights with a constant *c* > 1.

Any solution to Eq. S2 can, in theory, also be found with the optimization

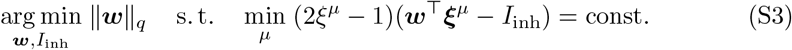

but this process would, in practice, be incompatible with a homeostatic process that keeps the weight norm fixed.

In machine learning terms, each neuron can be viewed as a linear classifier that discriminates *M* random input patterns ***ξ***^*µ*^ according to the output labels *ξ*^*µ*^. In this context, solving Eq. S2 (or S3) is equivalent to maximizing the classification margin with respect to the *L*_*q*_-norm, that is

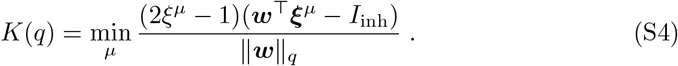

Given a fixed load *α* = *M/N* and pattern sparseness *f*, the maximum margin *K*^*∗*^ that a linear classifier can achieve is determined by a function *K*^*∗*^ (*α, f, q*). This defines the state of *optimal storage*, independently of the scaling of the weight vector ***w***. Historically, however, it is more common to rearrange the max-margin function so that the state of optimality instead is defined as the maximum load *α*^*∗*^(*K, f, q*) that can be attained with a fixed margin *K*. In this context, the largest possible storage load, at any margin, is referred to as the *critical capacity α* _*c*_, which is given by

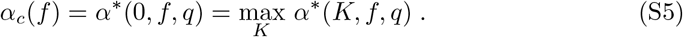

The reason this is independent of *q* is that a max-margin classifier at saturation (*α*^*∗*^ *→ α*_*c*_) has a vanishing margin (*K →* 0) and therefore no degrees of freedom to move. Hence, only a single solution exists at *α*_*c*_, regardless of which norm that is used to measure the margin.

As a notational rule, we use an asterisk (*) to denote any variable or function that is at optimal storage. Based on the two formulations of optimality, it is now possible to define the notion of *optimal learning* in two different ways:

#### S.1.1. Robustness maximization

By considering the state of optimal storage to be determined by *K*^*∗*^(*α, f, q*), we can define optimal learning as the process of finding the network configuration

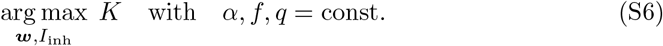

This is known as the *max-margin classifier* or *support vector machine* [88], and it is equivalent to our definition of consolidation in Eq. S2 (see Suppl. Fig. S8, dark arrows). The advantages of this approach are two-fold: First, it allows the network to flexibly attain maximal robustness and to operate optimally, without risk of catastrophic forgetting, at every storage load that is below critical capacity (i.e., *α* < *α* _*c*_). Second, it allows for this process to be carried out by an iterative learning rule that includes a homeostatic constraint on the weights.

### S1.2 Storage maximization

If one considers optimal storage to be defined by *α* ^*∗*^(*K, f, q*), it is natural to formulate optimal learning as the process of finding

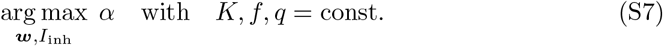

We refer to this as the *storage problem*. The main advantage of this approach is that it, in certain cases, is analytically tractable and allows for the optimal state of the network to be described with closed-form solutions in the mean-field limit *N →*∞. Here, we focus on three specific cases:

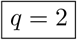

This solution is provided by Gardner [11] and is obtained under the weight scaling 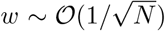 (see Suppl. Fig. S8a, light arrow). We use this to compute the maximum SNR with respect to neural noise. For technical details, see Supplementary Note S.2.1.

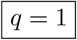

This solution can be found in references 12, 13, 16 and is obtained by keeping *I*_inh_ fixed and scaling the weights as *w ∼* 𝒪 (1/*N*) (see Suppl. Fig. S8b, light arrow). We use this solution to compute the maximum SNR with respect to synaptic noise in networks with two-factor synapses. For technical details, consult Supplementary Note S.2.2.

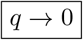

This solution is derived by Bouten et al. [82] by optimally diluting Gardner’s solution for *q* = 2. It describes the maximum amount of pruning that can be supported by a network. For technical details, see Supplementary Note S.2.3.

As a model of memory consolidation, however, the mean-field formulation has a number of disadvantages. Mainly, it is unclear how to translate it to a biologically realistic iterative learning rule, given that the margin *K* is a constant that has to be fine-tuned, *a priori*, to the particular load *α* that the network needs to store. One way to avoid this issue is to assume that K always stays fixed and is hard-coded into the learning rule. This is the approach taken in references 13, 45, 62 and in our control model. However, as suggested in Figure 4, this type of learning can only achieve optimal storage once the network has accumulated enough patterns to match the fixed margin. Until this point in time, the network operates at suboptimal storage. Moreover, after optimal storage load has been reached, the network must maintain a steady-state of stored patterns, in order to avoid catastrophic forgetting [14].

Finally, we argue that the basic assumption that neural circuits have a fixed robustness and learn to maximize the amount of memories is problematic from an ethological perspective. It implies that the brain does not adapt to environmental cognitive pressures, but instead passively incorporates information as it is encountered, without allowing for further improvement in the encoding.

#### S.1.3. Geometrical interpretation

Instead of solving Eq. S2 directly in terms of ***w***, we derive our consolidation algorithm by maximizing the SNR in the space of sub-synaptic *u*-variables, by solving

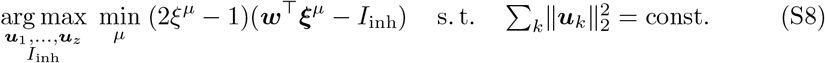

where the weight vector is composed of the Hadamard product ***w*** = ***u***_1_ *⊙ …⊙* ***u***_*z*_. At optimality, all sub-synaptic vectors align, so that ***u***_1_ = *…* = ***u***_*z*_ = ***u***. We prove this in the following theorem.

##### Theorem 1.

*Let 𝒬 be homogeneous objective function that obeys 𝒬*(cw) = c*𝒬*(w) *∀*c > 0, *where* w ≥0, *and consider the optimization problem*

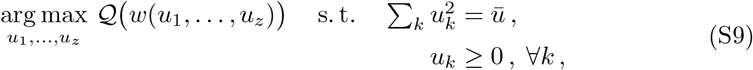

*where ū is a constant and w is parameterized as*

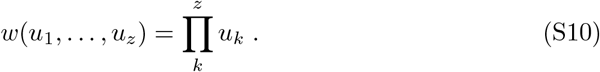

*Then, any local maximum* 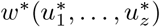 *must satisfy*

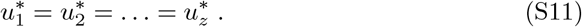

***Proof***. Consider a z-dimensional space spanned by the all the u-variables. In the positive orthant, the local maximum 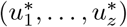 forms a rectangle together with the coordinate axes. This rectangle has volume *w*^*∗*^ and a diagonal of length 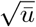. Recall that a rectangle with fixed volume minimizes the length of its diagonal only when all sides have equal length (equivalently, a rectangle with fixed diagonal length achieves maximal volume only when all sides are equal). In our case, this implies that for any candidate solution *w*^*∗*^ with unequal *u*^*∗*^-variables, a better solution can always be found with the following two steps:

Equalize all *u*^*∗*^-variables and generate a new solution *v*^*∗*^ with the same volume

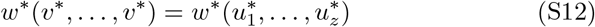

but with a shorter diagonal length

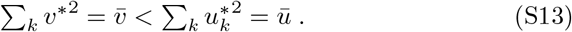

Rescale *v*^*∗*^ with the factor 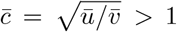 so that 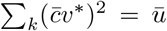, which satisfies the optimization constraint. The objective function now assumes the value

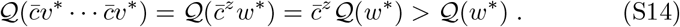

Thus, the new solution 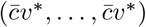 is superior. This proves that the candidate 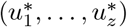 can never be an optimum, and that Eq. S11 therefore is a necessary condition. This argument can be generalized to *N* dimensions, where w is replaced by the vector ***w*** = (*w*_1_,…, *w*_*N*_). In this case, the two-step procedure is applied to each element of the vector separately.

This result allows us to rewrite Eq. S8 as

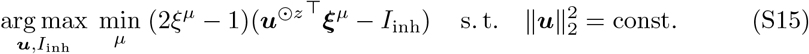

which, following a variable change ***u*** = ***w***^1*/z*^, is equivalent to

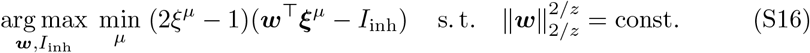

In other words, optimizing the SNR in u-space, using *z* components per weight, results in a weight vector that solves the original problem in Eq. S2 with exponent *q* = 2/*z*. In general, this type of regularized optimization yields progressively sparser solutions as *z* increases (i.e., *q* decreases). We provide an intuitive explanation for this phenomenon by analyzing the geometry of Eq. S2 from two perspectives: the *neural state space* and the *loss landscape*.

#### Neural state space

We consider a network of three neurons, and we study the two specific cases *z* = 1 and *z* = 2, which are equivalent to solving Eq. S2 with *q* = 2 and *q* = 1, respectively.

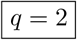

The solution to Eq. S2 is equivalent to a sign-constrained linear classifier at maximum margin *K*^*∗*^(*q* = 2). In Supplementary Figure S9a, we illustrate this solution in the two-dimensional state space of the afferent neural activity. Here, the optimal weight vector ***w***^*∗*^ and inhibition 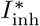 together define a classification boundary that correctly separates all patterns ***ξ***^*µ*^ and maximizes the Euclidean distance to the nearest items. The boundary is not biased towards any direction, so few entries in the normal vector are pushed to zero, which means that ***w***^*∗*^ is dense.

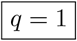

The solution to Eq. S2 is now equivalent to a sign-constrained linear classifier at maximum margin *K*^*∗*^(*q* = 1). We illustrate the state space representation of this solution in Supplementary Figure S9b. A theorem by Mangasarian [89] tells us that any classifier that maximizes the *L*_*q*_-margin, where *q* ≥ 1, corresponds, in geometrical terms, to a boundary that maximizes the 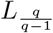 -distance to the nearest points. Consequently, at *K*^*∗*^(*q* = 1), the solution is a boundary that maximizes the *L*_*1*_-distance to all patterns ***ξ***^*µ*^. This forces the boundary to align with some of the coordinate axes, which zeros the corresponding weights and makes ***w***^*∗*^ sparse.

#### Loss landscape

We consider, as in the previous section, a neuron with two-dimensional input, and we define, as a simple example, the optimization problem

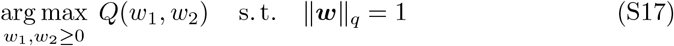

where *Q* is an objective function given by the paraboloid

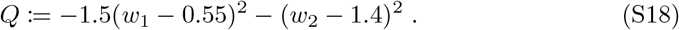

We present the *Q*-landscape, together with the constraint ‖***w*** ‖_*q*_ = 1, for different q-values, in Supplementary Figure S10a. As *q* is lowered, the shape of the constraint curve becomes more convex and moves the optimum closer to a sparse solution of type 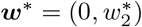.

We can numerically search for the optimum by performing projected gradient ascent (Suppl. Fig. S10b) according to the iterative algorithm

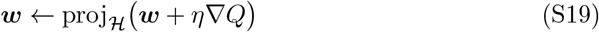

where *η* is the learning rate and the projection operator is defined as

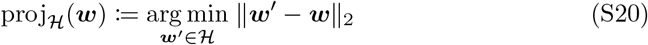

where the feasible set is given by : *H*= {***w***^*0*^≥0 : ‖***w***^*0*^‖_*q*_ = 1}. We analyze three specific cases of this process:

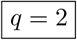 The projection operator is reduced to a multiplicative scaling, where

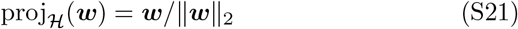

similarly to Oja’s rule [90]. This is compatible with the kind of homeostatic synaptic plasticity that has been observed experimentally [31], but is irreconcilable with the high degree of sparsity seen in cortical circuits, given that solutions generally are dense [71].

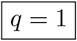 The projection operator is reduced to a subtractive adjustment, applied elementwise according to

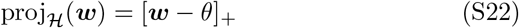

where [*·*] _+_ is the rectified linear function with a threshold θ that must be computed at every iteration, depending on ***w***, to satisfy ‖***w*** ‖_1_= 1. The resulting learning rule is now incompatible with biological homeostatic plasticity, but produces solutions with a sparsity comparable to cortical connectivity. Similar methods are used in references 16, 34, 72, 91.

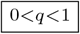 A closed-form expression for the projection operator is not available, as the shape of the constraint curve requires an anisotropic projection that, in general, adjusts weights by different amounts depending on ***w***. This poses a problem both from a modeling perspective and in terms of biological plausibility.

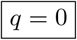 In this case, the projection operator is reduced to the the hard thresholding operation

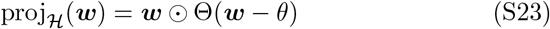

where Θ is the Heaviside function with a threshold *θ* that must be computed at every iteration, depending on ***w***, to satisfy ‖***w***‖_0_ = 1. For example, in the two-dimensional case, one chooses *θ* = min(*w*_1_, *w*_2_). This type of projection does not impose any form of homeostatic plasticity, and only prunes weights in order to produce solutions with a pre-defined level of sparsity. A similar method is used in reference 34.

We reconcile the need for multiplicative scaling with sparse solutions by expressing the weights as

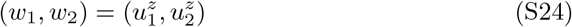

and solving

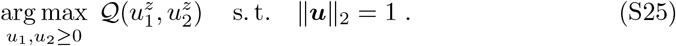

In Supplementary Figure S10c, we plot the reparameterized Q-landscape using, as an example, *z* = 3, together with the constraint curve ‖***w*** ‖_2*/*3_ = 1. The variable change deforms the landscape in such a way that the constraint curve can be reached with a multiplicative projection, even though the optimal solution remains sparse. The effect is the same for any pair of *z* and *q* = 2/*z*.

#### S.1.4. The gating function

Our derivation of the gating function *g* originates from the gradient calculation

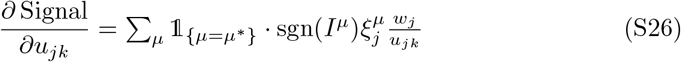

where we omit index *i* and use *µ*^*∗*^ = arg min*µ*| *I*^*µ*^ |. By replacing the indicator function with the Softmin, as in

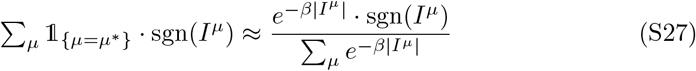

and then introducing

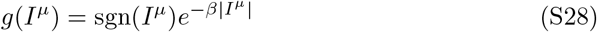

we arrive at the gradient approximation

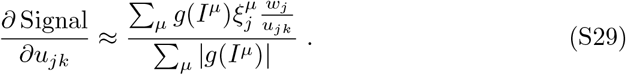

Note that the gating function takes on the shape of a *surrogate gradient* [92]. However, in contrast to the typical use-case of surrogate gradients, the performance of our model improves with higher *β*, as this reduces the discrepancy between the approximate and true gradient (Suppl. Fig. S11). In order to guarantee that the approximate gradient converges to the true gradient in the limit *β2192* ∞, the tails of *g* must decay to zero at a rate that is, at least, faster than a polynomial. We state this in the following theorem.

##### Theorem 2.

*Consider a general Softmin function*

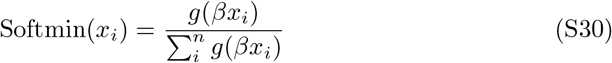

*where x*_*i*_ > 0 *∀i, β* > 0 *is an inverse temperature, and g*(x) *is a finite and strictly positive function that decays monotonically to zero as x →*∞. *Then*,

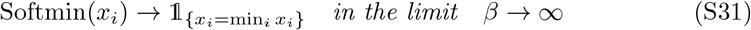

*iff g decays faster than a polynomial, that is, g(x) ∼o(x^−c^), with 0 < c <*.

***Proof***. Let *x*_1_ and *x*_2_ denote the smallest and second smallest *x*_*i*_. The convergence in Eq. S31 is equivalent to

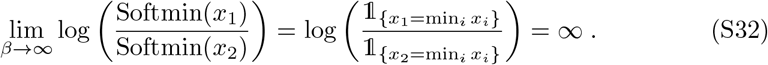

At the same time, we have

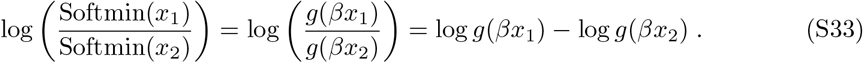

We combine Eq. S32 with S33 and obtain

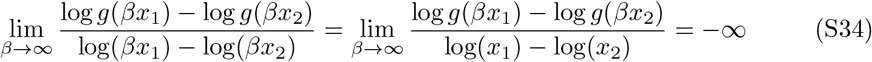

where the minus sign on the right-hand side is due to *x*_1_ < *x*_2_. This condition must hold for any pair of *x*_1_ and *x*_2_, no matter how close they are to each other. In the limit *x*_2_ → *x*_1_, Eq. S34 is equivalent to

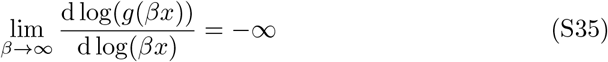

which states that the slope of g, in logarithmic space, cannot be bounded, but must tend to −∞. In other words, the tail of g must decay faster than a line in logarithmic space, and, thus, faster than a polynomial in linear space, which means 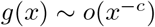.

#### S.1.5. The homeostatic function

As shown in the Methods, the consolidation model with two-factor synapses can be expressed in continuous time with the differential equation

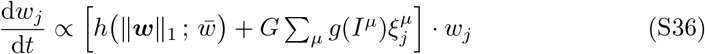

where we omit index i. The dynamics of the homeostatic term is determined by the function h, which is defined as 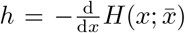, where *H* represents a homeostatic penalty function that is zero at 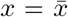 and increases monotonically everywhere else. This can be viewed as a generalized formulation of homeostatic plasticity, which, depending on the exact shape of H, can be reduced to specific instances of plasticity models that have been proposed in previous work. Consider the following three cases:

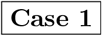 If we choose the penalty function to be

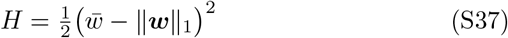

we obtain the homeostatic function

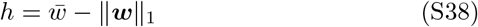

which is identical to the homeostatic scaling rule introduced by Renart et al. [35], albeit expressed in terms of the summed weights instead of input firing rates.

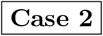 If we instead define the penalty as

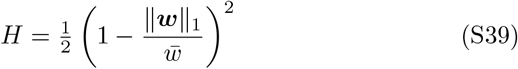

we retrieve the homeostatic function

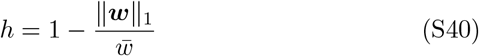

which is the homeostatic rule introduced in by Toyoizumi et al. [36], but expressed in terms of summed weights instead of the input currents.

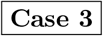 A third alternative for the penalty function is

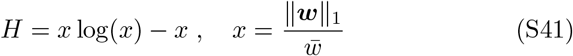

which yields the homeostatic function

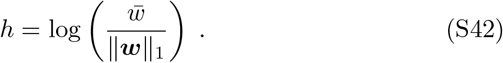

This type of homeostatic scaling has, to the best of our knowledge, not been proposed previously in the literature.

It is important to note that even though all homeostatic rules regulate the average synaptic weight, they do so by monitoring different quantities. In case 1, the rule depends on a raw deviation from the set-point, while, in case 2, it depends on the percentage of the deviation. In the third case, the homeostatic rule depends only on the ratio of ‖***w***‖_1_ relative the set-point.

#### S.1.6. Related algorithms

In this section, we explain the link between the consolidation model and other iterative learning algorithms. The expression for Δ*u*_*ijk*_ in Eq. 47 can be seen as a generalized weight update rule, which, depending on the value of *β*_*i*_, can be reduced to three well-known algorithms from the machine learning literature:

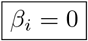 In this case, our update rule is reduced to the conventional gradient ascent procedure

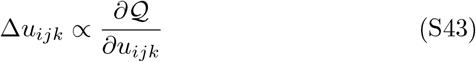

where the objective function 𝒬 is the average signal across all patterns, given by

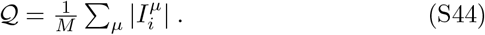

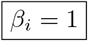 This case is equivalent to the *normalized* gradient ascent algorithm [80]

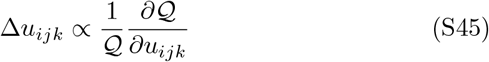

applied to the exponential objective function

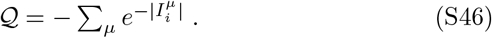

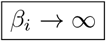 In this limit, our update rule becomes identical to the *batch perceptron* algorithm [81]

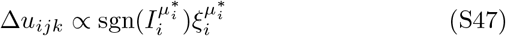

where 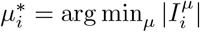.

Both the normalized gradient and the batch perceptron were originally introduced as margin-maximizing learning rules. Indeed, as we demonstrate in Supplementary Figure S11, the performance of our algorithm improves with increasing /3_*i*_. At very high /3_*i*_, it appears to converge to the batch perceptron, which consistently performs best.

### S.2. Theoretical solutions

#### S.2.1. Maximal neural noise robustness

To calculate the theoretically highest possible SNR with respect to neural noise, we use the solution for the maximum margin *K*^*∗*^ (*α, f, q* = 2), which we obtain using the maximum load *α*^*∗*^(*K, f, q* = 2) and solving for *K*. The maximum load *α*^*∗*^ is the solution to Eq. S7 with *q* = 2, and is provided by Gardner [11] in the form

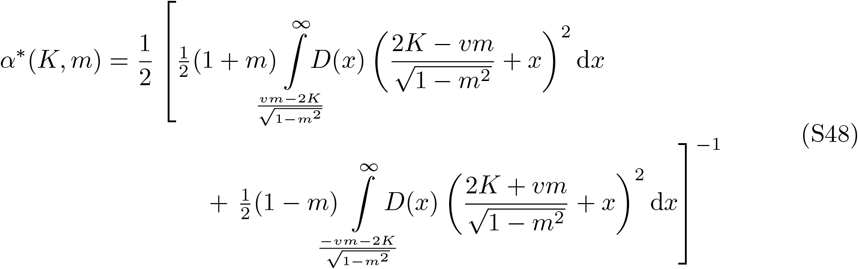

where *v* is given by the solution to the equation

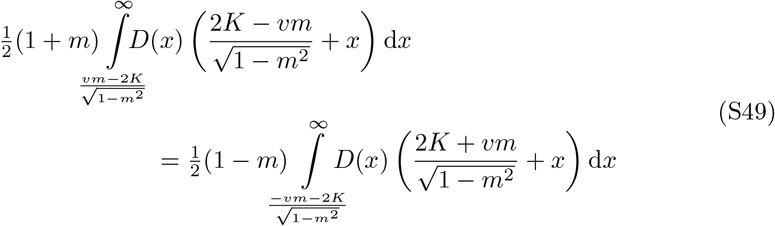

and *D* is the standard normal distribution

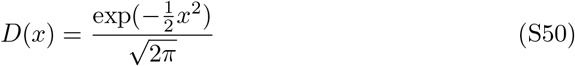

and *m* is the pattern magnetization, which simply reflects the activity level *f* according to

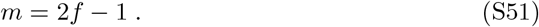

In the specific case of balanced patterns (*f* = 0.5), Eq. S48 is evaluated at m = 0 and reduced to

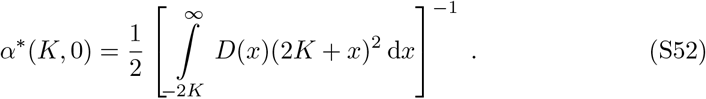

Note that both *α*, ^*∗*^ and K have been adjusted with a factor 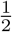 relative the original solution by Gardner. This accounts for the fact that we allow only non-negative weights [93] and use pattern-values in {0, 1}, while the original solution was derived for unconstrained weights and patterns in *{±*1*}*. The SNR is now computed as

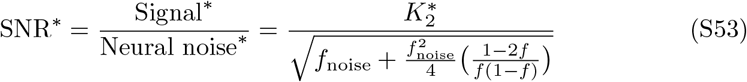

where 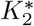 is shorthand for 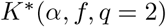.

#### S.2.2. Maximal synaptic noise robustness

##### Single-factor synapses

In the case of *z* = 1, synaptic noise depends on the fraction of non-pruned weights *f*_*w*_ (see Methods). In order to compute the highest possible SNR with respect to synaptic noise, it is therefore necessary to derive the fraction of weights that a sign-constrained linear classifier exhibits at 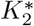. We denote this optimal fraction 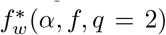. At activity level *f* = 0.5, this is, in fact, known to be exactly 50%, regardless of storage load [83]. Given a weight norm ‖***w***‖_2_, the optimal signal can, according to Eq. S4, be written as 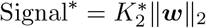, which gives us the maximal SNR

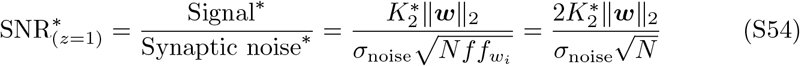

where the last equality is obtained by inserting 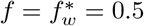.

##### Two-factor synapses

In the case of *z* = 2, we can compute the highest possible SNR with respect to synaptic noise using the solution for the maximum margin *K*^*∗*^(*α, f, q* = 1), which is obtained from the maximum load *α*^*∗*^(*K, f, q* = 1) after solving for *K*. The maximum load *α*^*∗*^ is the solution to Eq. S7 with *q* = 1. This was first published in references 12, 13. Here, however, we use the solution reported by Zhang et al. [16], which is expressed as

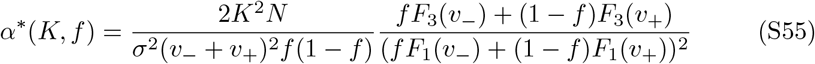

where the variables (*x, v*_*−*_, *v*_+_,*σ*) are given by the solution to the system of equations

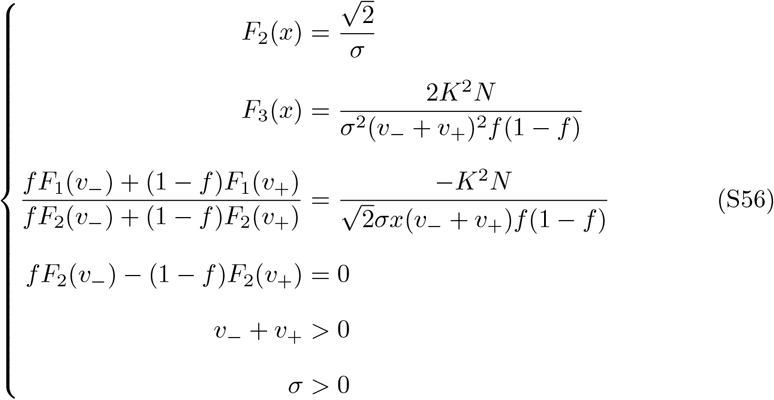

where we use the auxiliary functions

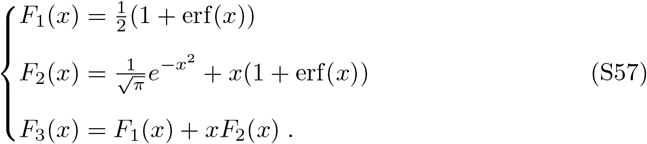

This is also used to compute the optimal weight fraction

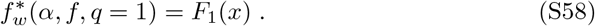

The maximal signal at a given weight norm ‖***w*** ‖_1_ is now given by 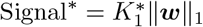 (see Eq. S4), which yields the maximal SNR

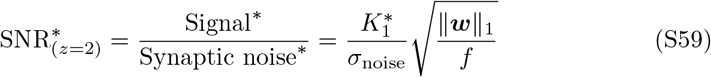

where 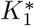 is shorthand for *K*^*∗*^ (*α, f, q* = 1).

#### S.2.3. Maximal pruning

In the limit *z* 2192 ∞, our definition of consolidation is equivalent to a maximization of the *L*_0_-margin, which, according to Eq. S3, can be formulated as a minimization of the number of observable weights ‖ ***w* ‖**_0_ relative the signal. This, in other words, describes the maximum fraction of weights that can be pruned by a neuron, without losing any of the stored patterns. In order to compute this, we turn to the optimal storage definition in Eq. S7 and supplement it with an additional constraint that requires the optimum to have a desired weight fraction *f*_*w*_. The result is a new storage optimization

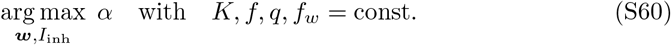

whose solution now is described by the maximal storage load *α*^*∗*^(*K, f, q, f*_*w*_). The value of *α*^*∗*^ that is attained at the smallest possible margin, that is *K* = 0, is the critical capacity

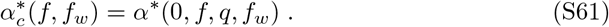

Note, again, that this function is independent of *q*, as the zero-margin solution is the same for all *q*. The highest degree of pruning is now determined by the lowest possible *f*_*w*_ at a given *α*_*c*_. We obtain this by solving for *f*_*w*_ in Eq. S61 and write it as the function

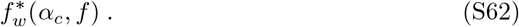

In the case of balanced patterns, *f* = 0.5, a derivation of 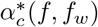 can be found in the work by Bouten et al. [82]. The result is

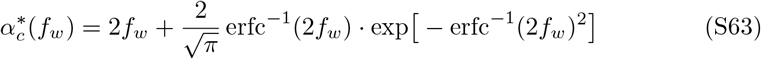

where 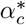 has been adjusted with a factor 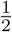 relative to the original solution in order to account for the sign-constrained weights [93]. We have also scaled 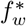 with a factor 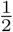 relative to the original solution. We motivate this with a symmetry argument: The original, unconstrained solution always has a weight distribution that is symmetric and centered at zero, with an equal number of positive and negative weights [82]. Intuitively, it is therefore reasonable to expect that a sign-constraint causes precisely half of all weights to have the wrong sign and to be pruned to zero. This has, indeed, been proven to be true at 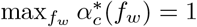 [83], where we have

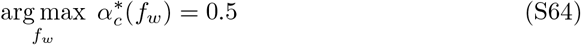

and we conjecture that the same applies for all ff_*c*_.

### S.3. Derivation of sparseness

In the main text, we define sparseness as

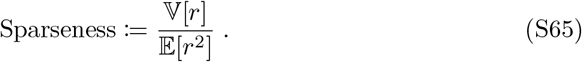

In practice, this metric is applied to a sample of neural stimulus responses, acquired either from simulations or biological experiments. In this case, we replace the variance and expectation with the *unbiased* sample estimates, so that

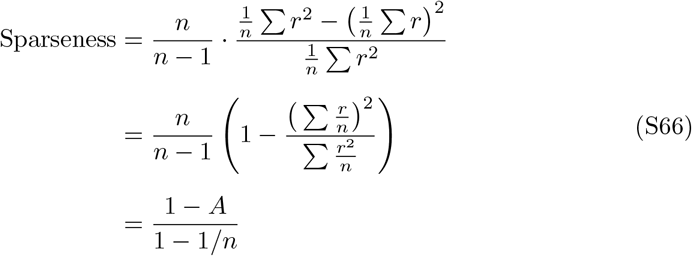

where *n* is the number of samples and

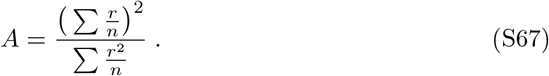

This is the way sparseness is formulated in the literature [85, 86].

### S.4. Extended synaptic noise analysis

As observed in the main text, datasets with smaller sample sizes and longer sampling intervals have a scaling exponent that generally increases for depression and decreases for potentiation (i.e., it diverges). This is particularly evident in the case of the longest sampling intervals (Δ*t* ≥48 h; Fig. 5c, third group of data) where the exponent is 0.38 ± 0.04 for potentiation, and higher than one (1.09±0.03) for depression, consistent with previous analyses of this type [70].

To further verify these observations, we artificially decrease the sampling frequency in each dataset by sub-sampling measurements across time. More specifically, instead of extracting all weight changes over the original sampling interval Δ*t*, we select only weight changes between two measurement separated by an interval Δ*t*_sub_ = *n ·* Δ*t*, where *n* = 2, 3,…, *n*_max_, and *n*_max_ *·* Δ*t* is the total length of the experiment. We then re-compute the scaling exponent as a function of the new sampling interval Δ*t*_sub_ (Suppl. Fig. S7). Results *within* datasets corroborate those *across* datasets: as the sampling interval increases, the exponent diverges from ∼0.6, by going above one for synaptic depression and decaying close to zero for potentiation. These same trend is found in the simulated data.

### S.5. Simulation parameters

**Table S1.**
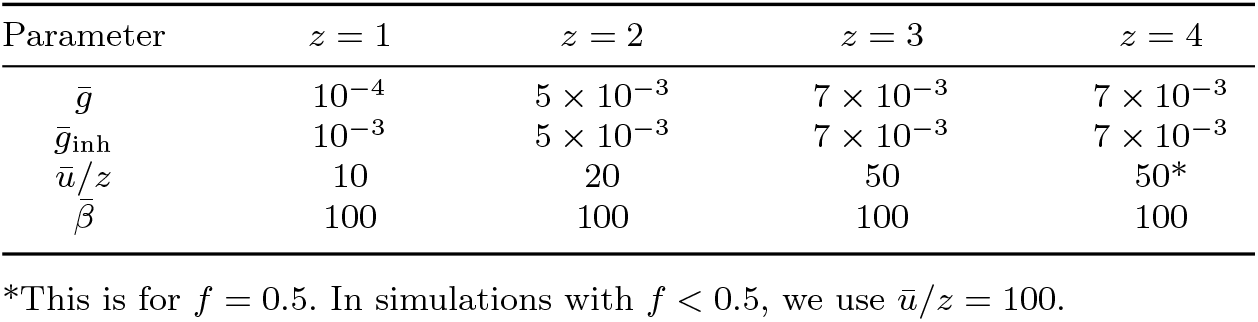
Simulation parameters for Figures 2 and 4.

**Table S2.**
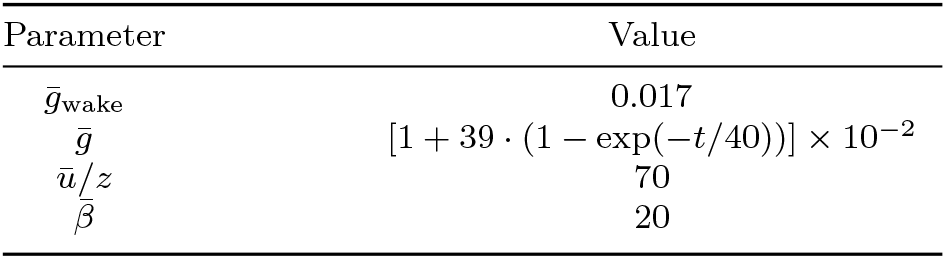
Simulation parameters for Figure 3. During sleep, the learning rate increases exponentially with a time constant of 40 replay cycles (*t* denotes the cycle).

**Table S3.** Simulation parameters for Figures 5 and 6.

| Parameter | Value |
| --- | --- |
| $\sigma_{\text{noise}}$ | 0.05 |
| $u_0$ | 0.1 |
| $\tau$ | 30 |
| $dt$ | 0.005 |
| $T_{\text{sample}}$ | 1 |
| $T_{\text{sim}}$ | 1000 |

### S.6. Metadata for synaptic imaging

The following three tables contain details about the experimental data used to produce Figures 5, 6, and S7.

**Table S4.**
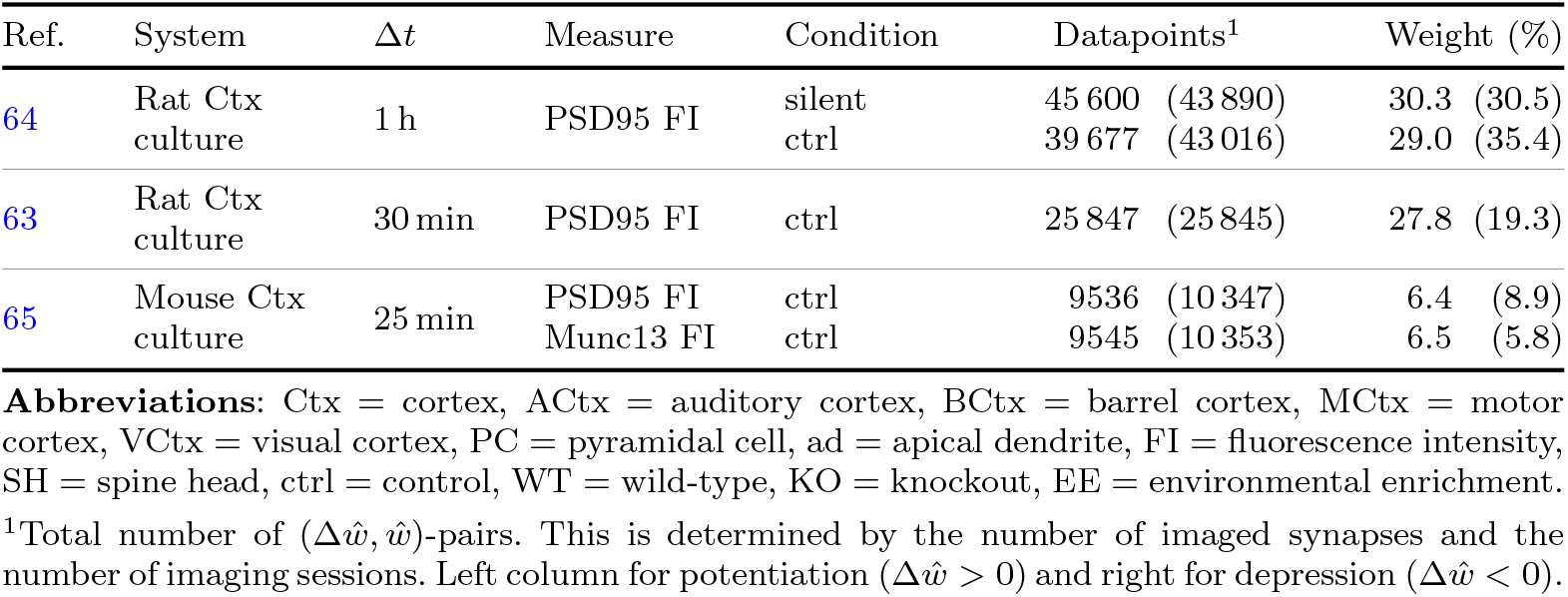
Description of synaptic data with large sample sizes and short sampling intervals.

**Table S5.** Description of synaptic data with small sample sizes and short to medium sampling intervals. Notation as in Table S4.

| Ref. | System | $\Delta t$ | Measure | Condition | Datapoints | Weight (%) |
| --- | --- | --- | --- | --- | --- | --- |
| 29 | Mouse MCtx L2/3 PC in vivo | 7 h | GluA1 FI | sleep | 1039 (1270) | 30.2 (28.2) |
|  |  |  |  | wake | 346 (405) | 9.8 (7.6) |
|  |  |  | SH FI | sleep | 1107 (1202) | 22.3 (19.5) |
|  |  |  |  | wake | 371 (380) | 7.7 (4.9) |
| 67 | Mouse VCtx L5 PC-ad in vivo | 10 min | SH FI | WT | 238 (237) | 3.8 (2.9) |
|  |  |  |  | Fmr1-KO | 714 (719) | 16.0 (17.1) |
| 69 | Mouse VCtx L5 PC-ad in vivo | 30 min | PSD95 area | EE | 169 (280) | 2.6 (8.6) |
|  |  |  |  | ctrl | 105 (228) | 1.4 (6.5) |
|  |  |  | SH area | EE | 237 (215) | 4.3 (2.4) |
|  |  |  |  | ctrl | 161 (169) | 1.9 (2.4) |

**Table S6.** Description of synaptic data with long sampling intervals. Notation as in Table S4.

| Ref. | System | $\Delta t$ | Measure | Condition | Datapoints | Weight (%) |
| --- | --- | --- | --- | --- | --- | --- |
| 66<sup>1</sup> | Mouse BCtx L2/3/5 in vivo | 96 h | Bouton FI | ctrl | 12 829 (12 773) | 72.0 (57.3) |
| 52 | Mouse ACtx L5 PC-ad in vivo | 96 h | SH FI | ctrl | 2459 (2552) | 16.5 (31.3) |
| 67 | Mouse VCtx L5 PC-ad in vivo | 48 h | SH FI | WT | 350 (404) | 4.7 (4.2) |
|  |  |  |  | Fmr1-KO | 417 (461) | 6.0 (6.4) |
| 68 | Mouse MCtx L5 PC-ad in vivo | 72-96 h | SH area | ctrl | 168 (244) | 0.8 (0.8) |

<sup>1</sup>We included only measurements for which the bouton detection probability was $> 90\%$ .

### S.7. Supplementary figures

**Fig. S1.**
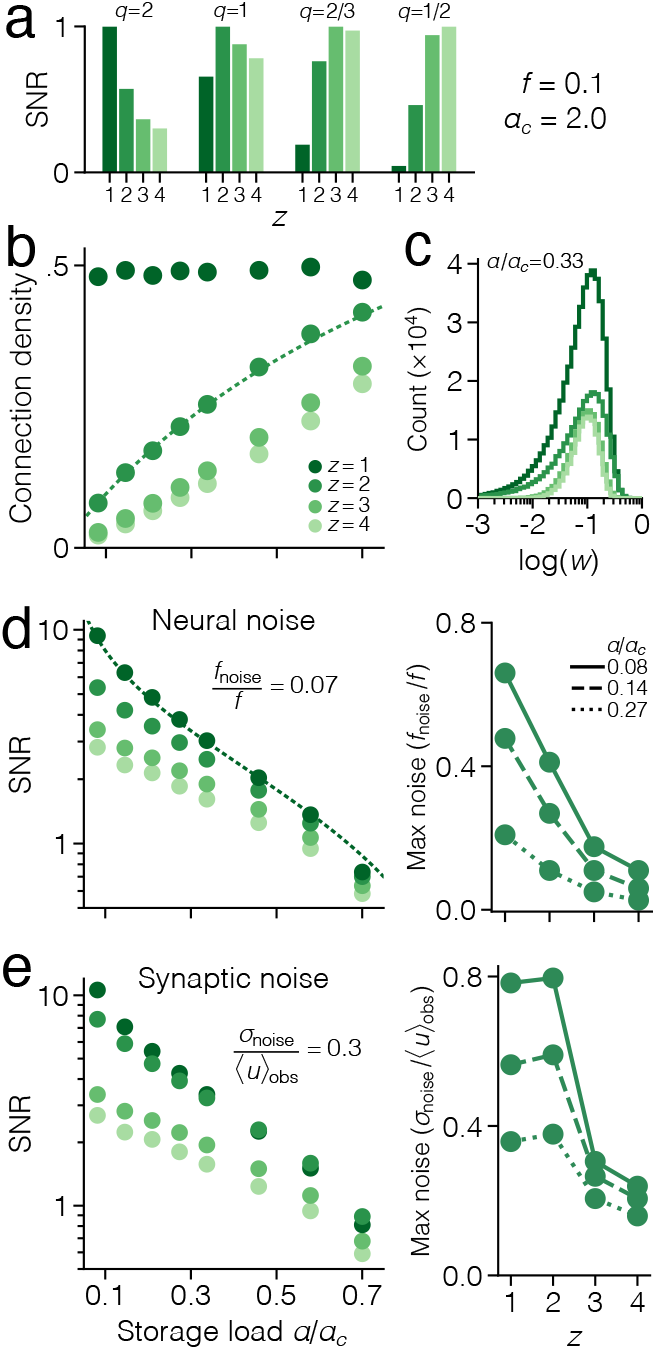
Simulated consolidation with low-activity patterns. The same type of results as Figure 2 but with *f* = 0.1. **(a)** SNR with respect to noise scaling *q*, at *α/α*_*c*_ = 0.08 (mean over 10^3^ neurons). Weights are normalized to 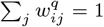 and the maximal SNR, for a given *q*, is scaled to one. Connection density. Dashed line corresponds to theory for *z* = 2. **(c)** Distribution of weights (mean scaled to 10^*-*1^). **(d)** SNR with respect to neural noise (*q* = 2; left) and highest level of tolerated neural noise in tests of pattern recall (right). Dashed line corresponds to theory for *z* = 1. **(e)** SNR with respect to synaptic noise (*q* = 2−2*/z*; left) and highest level of tolerated synaptic noise in tests of pattern recall (right).

**Fig. S2.**
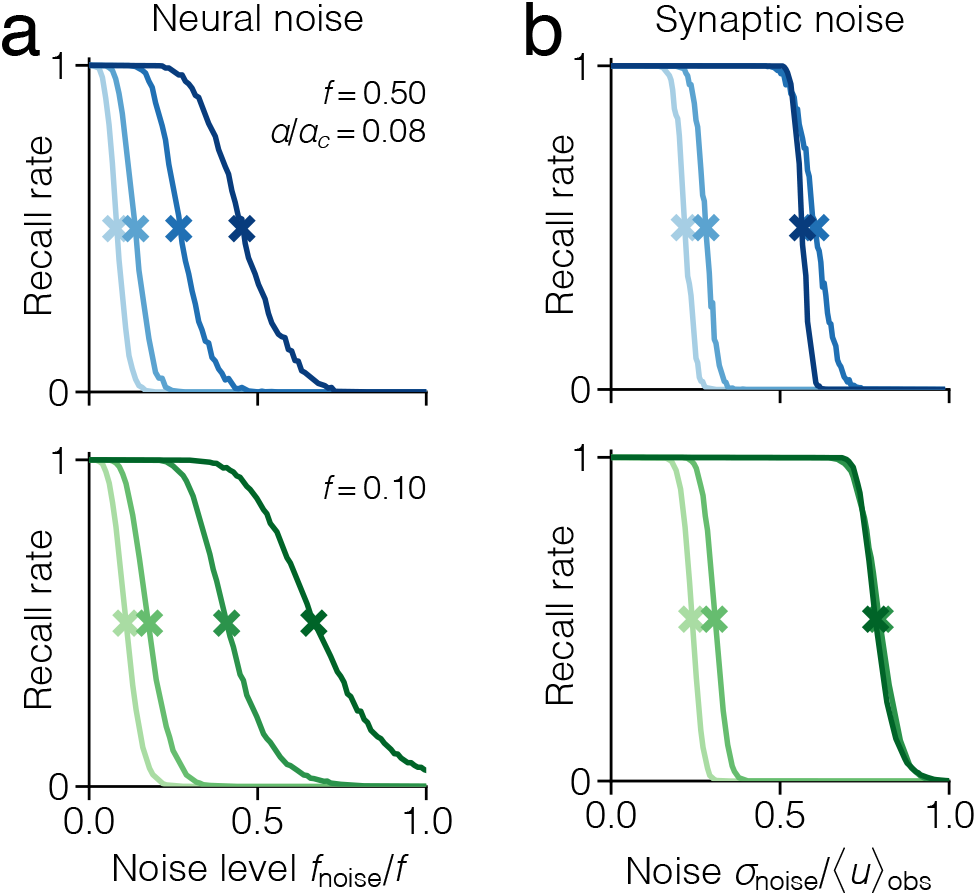
Empirical robustness evaluation. The fraction of memories that can be successfully retrieved (i.e., recall rate) as a function of **(a)** neural noise and **(b)** synaptic noise, in networks with pattern activity levels *f* = 0.5 (blues) and *f* = 0.1 (greens). Crosses indicate where the recall rate falls below 50%. This defines the highest level of tolerated noise.

**Fig. S3.**
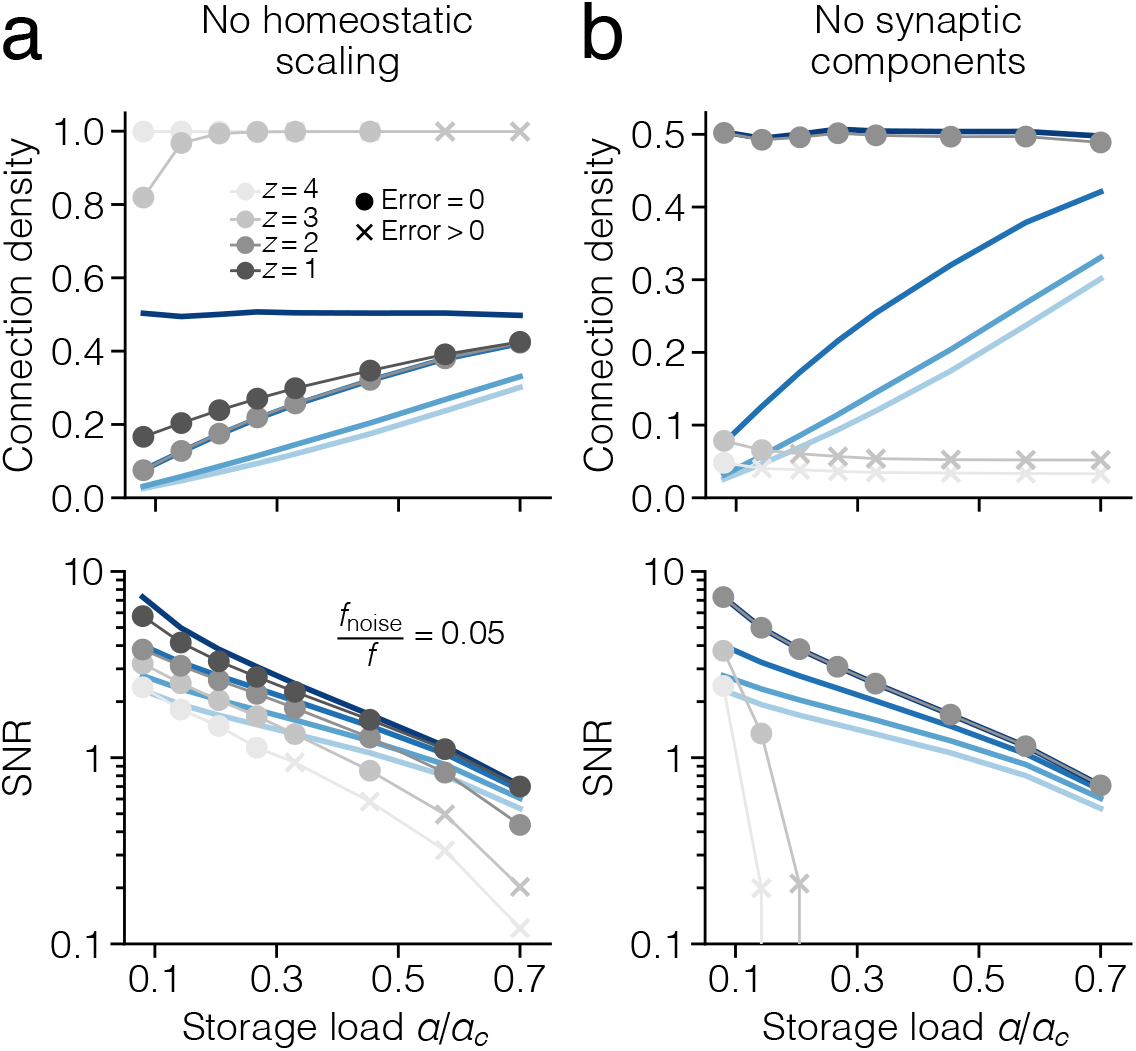
Ablated consolidation model. Connection density (top) and SNR with respect to neural noise (bottom) after consolidation with ablated (gray) and intact (blue) consolidation model. All markers correspond to means over 10^3^ neurons, but circles indicate cases where the network manages to find a solution with *E* = 0 in 2 ×10^6^ replay cycles, while crosses indicate cases where the network fails to find such solutions. **(a)** Consolidation without homeostatic scaling. With the exception of *z* = 2, the network fails to converge to any meaningful results. Simulation parameters as in Figure 2. **(b)** Consolidation with homeostatic scaling but only single-factor synapses (i.e., *z* = 1). Due to the multiplicative projected gradient ascent, the solution either coincides with the intact *z* = 1 solution, or, once again, fails to converge to anything meaningful. Simulation parameters as in Figure 2, but with learning rates 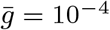 and 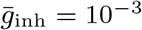.

**Fig. S4.**
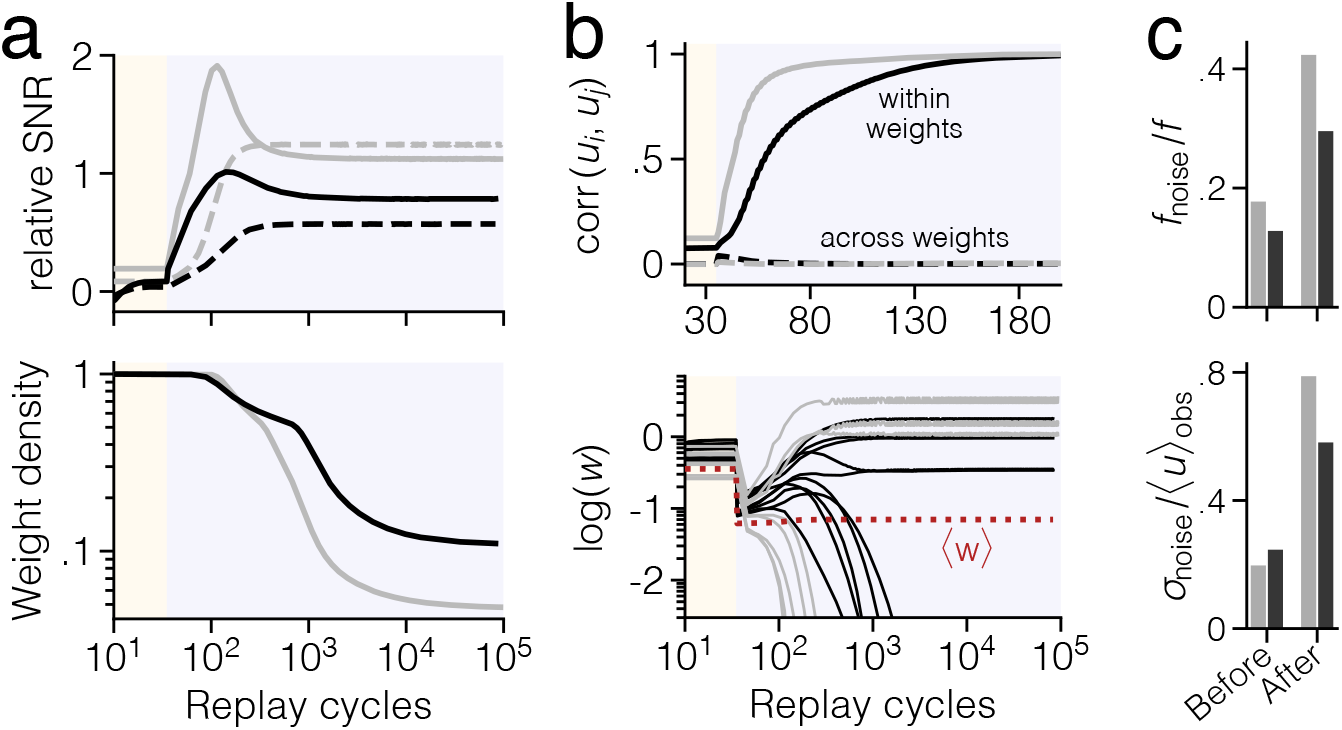
Memory formation and stabilization in wakefulness and sleep. Simulation of wakefulness (yellow background) and sleep (violet background) with high load (black; *α*= 0.44) and low load (gray; *α* = 0.2). **(a)** Relative SNR (top) and weight density (bottom) over replay cycles. Solid curves represent neural noise (*q* = 2) and dashed curves synaptic noise (*q* = 1). Scaling of the SNR-axis is arbitrary. **(b)** Top panel shows the pairwise Pearson correlation between subsynaptic components *u*_*ijk*_ within the same weight (same *j*, different *k*) and across different weights (same *k*, different *j*). Bottom panel shows the weight trace for a subset of synapses. **(c)** Maximum tolerated neural and synaptic noise before and after sleep.

**Fig. S5.**
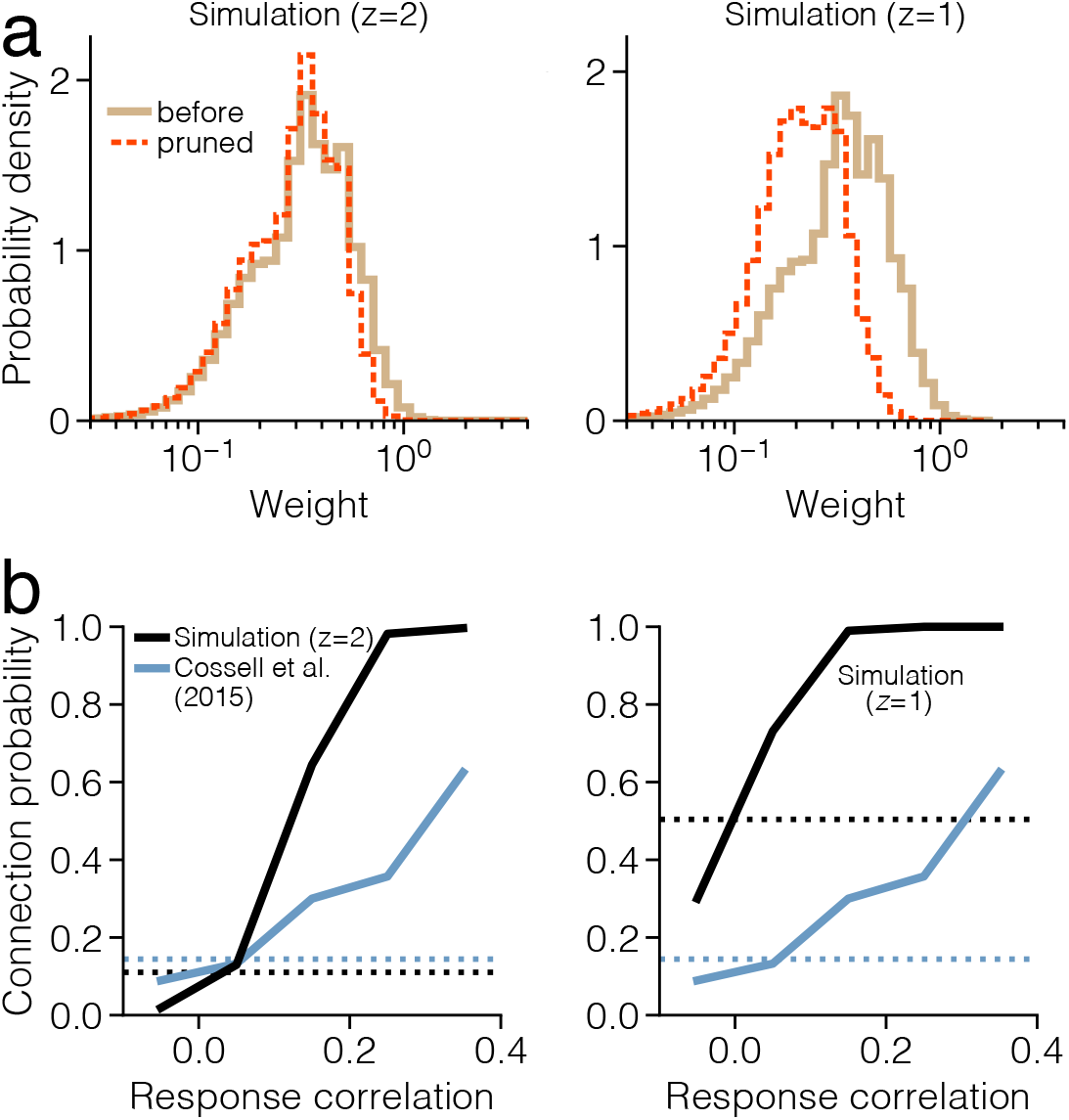
Comparison of dense and sparse consolidation. Simulation of wakefulness (few-shot learning) and sleep (consolidation) in a network with *z* = 2 (sparse; left) and *z* = 1 (dense; right), with low-activity patterns *f* = 0.05 at *α* = 0.44. **(a)** Distribution of pre-sleep and pruned weights. The degree of pruning is lower in the dense case than in the sparse case. Consequently, the distribution of pruned weights no longer overlaps with the distribution of pre-sleep weights. **(b)** Connection probability as a function of response correlation. The dense network, which always converges to solutions with roughly 50% connection probability, cannot reproduce the low level of connection probability observed in rodent visual cortex [53] (blue).

**Fig. S6.**
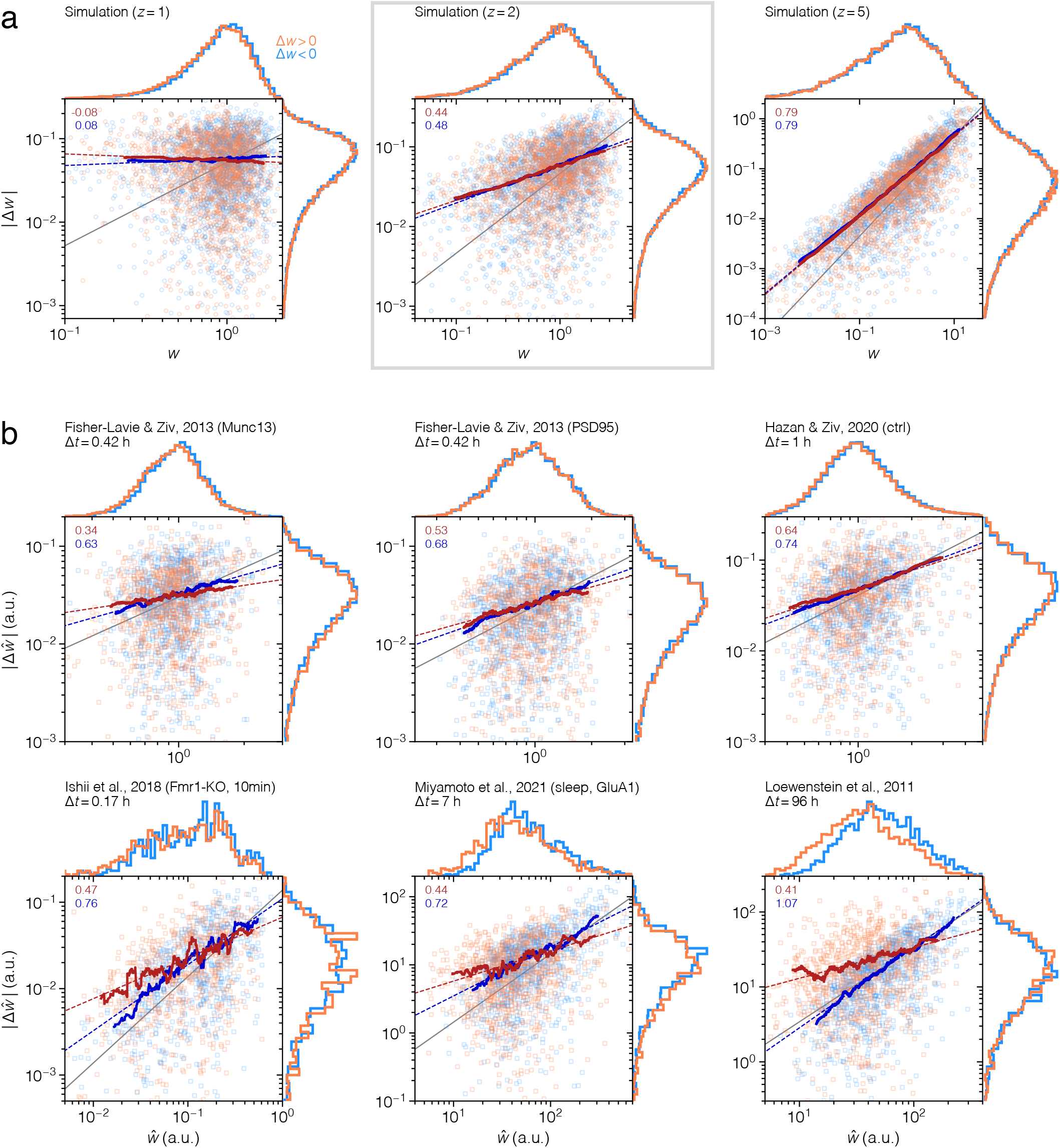
Extended synaptic fluctuation data. **(a)** Absolute weight change as a function of initial weight in simulated data with *z* = 1 (left), *z* = 2 (middle), and *z* = 5 (right), for potentiation (orange) and depression (blue). Solid lines are moving averages, and dashed lines are linear fits to the solid lines (slope value shown in upper left corner). The straight solid lines suggest a power-law in the original data, and their slope (i.e., the power-law exponent) approximately obeys the scaling law *q* = 1−1*/z*. The identity line (gray) has slope 1, and is included for comparison. **(b)** The same type of plot as in **a**, but for experimental measurements of dendritic spine sizes in cortical neurons, across different datasets. The sampling time is denoted with b.*t*.

**Fig. S7.**
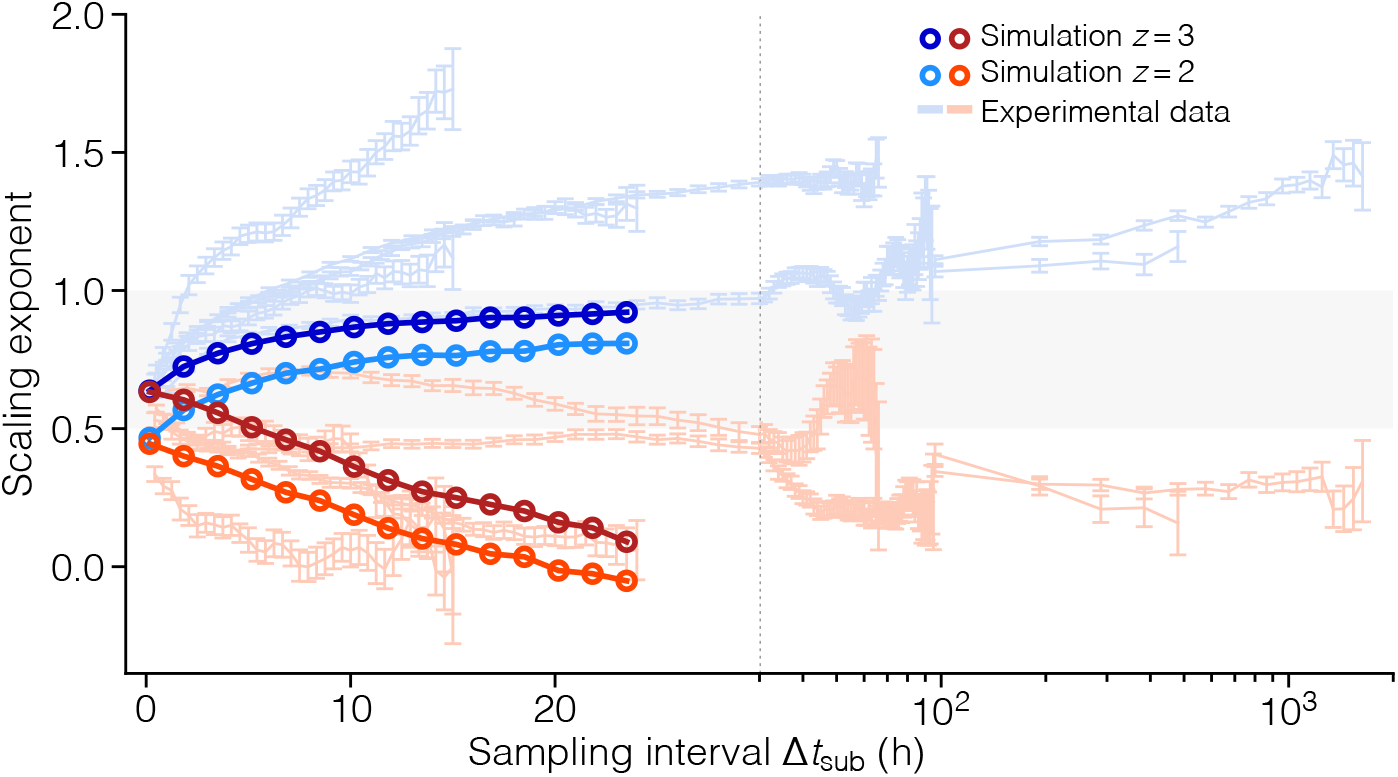
Synaptic noise scaling in subsampled data. The scaling exponent of synaptic fluctuations as a function of the sampling interval b.*t*_sub_, in simulated and experimental data [52, 63–66] (mean ±SE, estimated as in Fig. 5). The sampling interval is artificially lengthened by sub-sampling datapoints across time. The scaling exponent generally diverges by increasing for depression (blue markers) and decreasing for potentiation (orange markers).

**Fig. S8.**
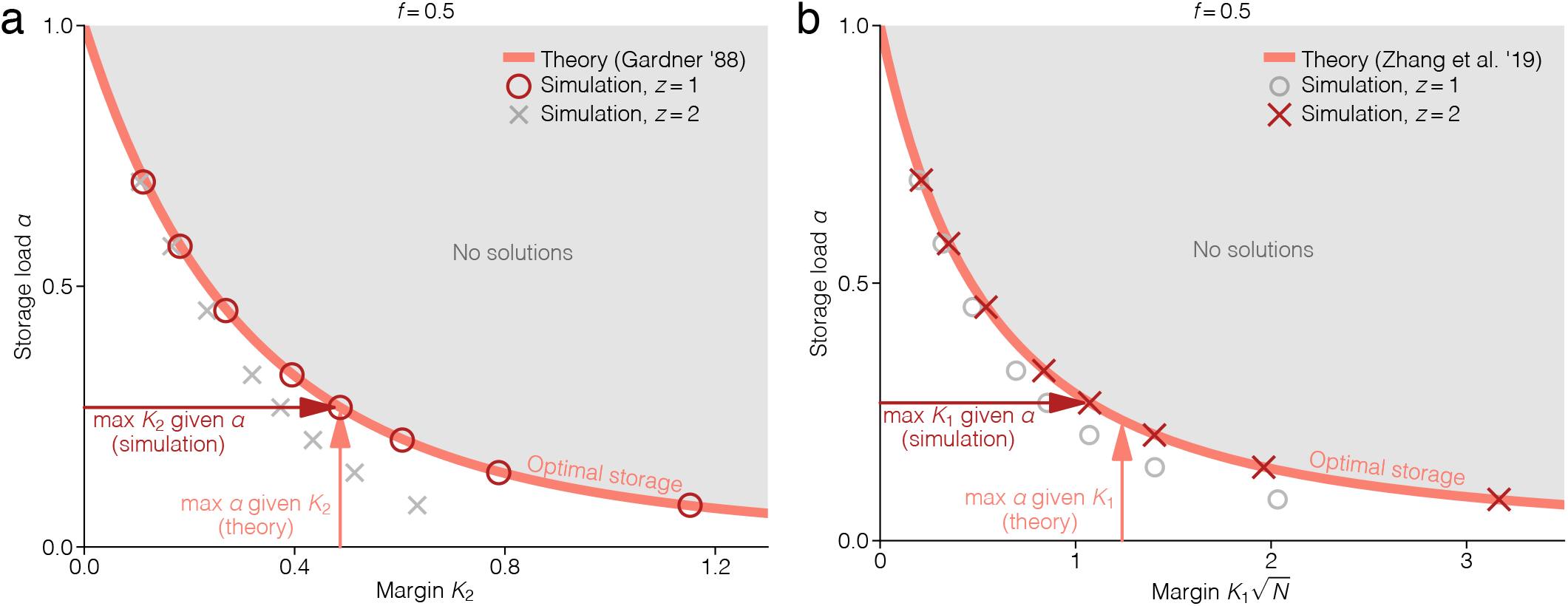
Comparison of the max-margin and max-storage formalisms. **(a)** Storage load *α*as a function of margin *K*_2_, shorthand for *K*(*q* = 2). The optimal storage curve is the function *α*^*∗*^(*K, f, q*) with *f* = 0.5 and *q* = 2, as reported by Gardner [11]. This is obtained by solving the *storage problem* in Eq. S7, where *α* is maximized, given a fixed margin (pink arrow) in the mean-field limit. The same optimal storage configuration can also be found by solving the corresponding *max-margin problem* in Eq. S6, where *K* instead is maximized, given a fixed load (brown arrow). This is what our consolidation model is derived to do. Indeed, it retrieves the solution when *z* = 1, as this maximizes *K*_2_, but not when *z* = 2, as this maximizes *K*_1_. **(b)** Storage load *α*as a function of margin *K*_1_, shorthand for *K*(*q* = 1). The optimal storage curve is the function *α*^*∗*^(*K, f, q*) with *f* = 0.5 and *q* = 1, as formulated by Zhang et al. [16]. Using our consolidation model, we now find the optimal storage solution when *z* = 2, as this maximizes *K*_1_, but not when *z* = 1, as this maximizes *K*_2_. The simulation results are the same as in Figure 2.

**Fig. S9.**
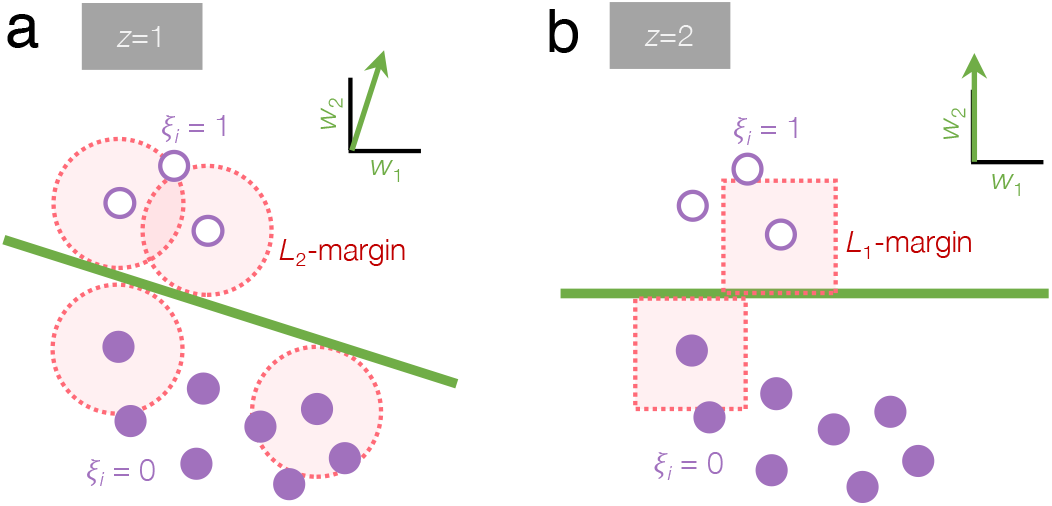
Dense and sparse consolidation in neural state space. In a network of *N* = 3 neurons, we consider a single neuron *i* and observe the state space of its two neighboring neurons. The weight vector ***w***_*i*_ = (*w*_1_, *w*_2_) and the inhibition *I*_inh,*i*_ define a linear classification boundary (green) that separate all patterns (circles) according to the labels *ξ*_*i*_ = 1 (white) and *ξ*_*i*_ = 0 (purple). For the sake of simplifying the illustration, we use real-valued patterns, but the same argument holds for the binary case. **(a)** Consolidation with *z* = 1 is equivalent to a maximization of the *L*_2_-margin, which means that the *L*_2_-distance between the boundary and the nearest patterns is maximized (red circles). The solution is typically dense, which means that 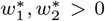 Consolidation with *z* = 2 is equivalent to a maximization of the *L*_1_-margin, which means that the *L*_*1*_-distance between the boundary and the nearest patterns is maximized [89] (red squares). The boundary is now forced to align with the one of the coordinate axes, thus rendering the solution sparse, such that 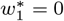.and 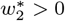.

**Fig. S10.**
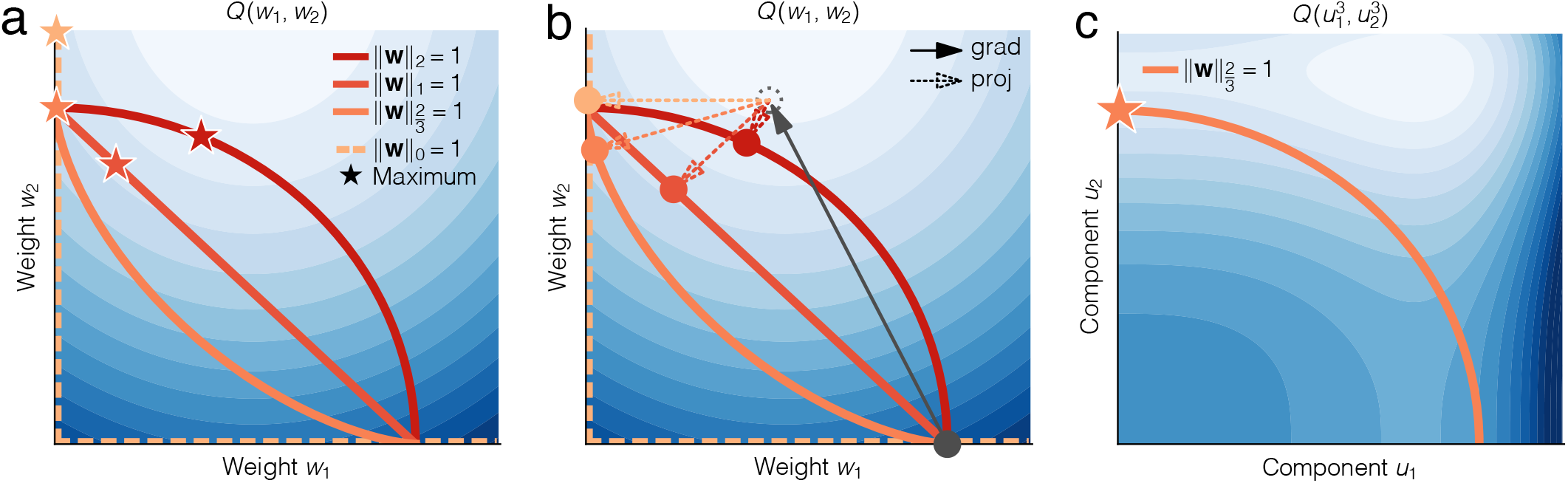
Dense and sparse consolidation in the loss landscape. **(a)** The landscape of the objective function *Q*(*w*_1_, *w*_2_) (blue; lighter hues closer to max) together with the feasible set under constraints of type ‖***w***‖_*q*_ = 1 (orange curves). In general, a lower *q* pushes the optimal weight vector (star) closer to a sparse configuration, in which 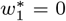 and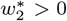. Indeed, for 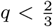, the solution is sparse. **(b)** Projected gradient descent in the *Q* -landscape involves first a gradient step (solid arrow), followed by a projection to the feasible set (dashed arrow). The projection can be multiplicative (*q* = 2), additive (*q* = 1), or a hard thresholding (*q* = 0). However, for fractional norms (0 *<q <* 1), the projection is generally anisotropic, which means that weights are adjusted by different amounts, depending on their relative size to each other. **(c)** We can make the projection to any fractional norm curve multiplicative, by performing the optimization in the re-parameterized landscape 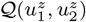 if we choose the number of components *z* = 2*/q*. For example, projections to 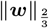 (orange curve) become multiplicative with *z* = 3. The optimum remains sparse, with 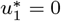 and 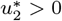.

**Fig. S11.**
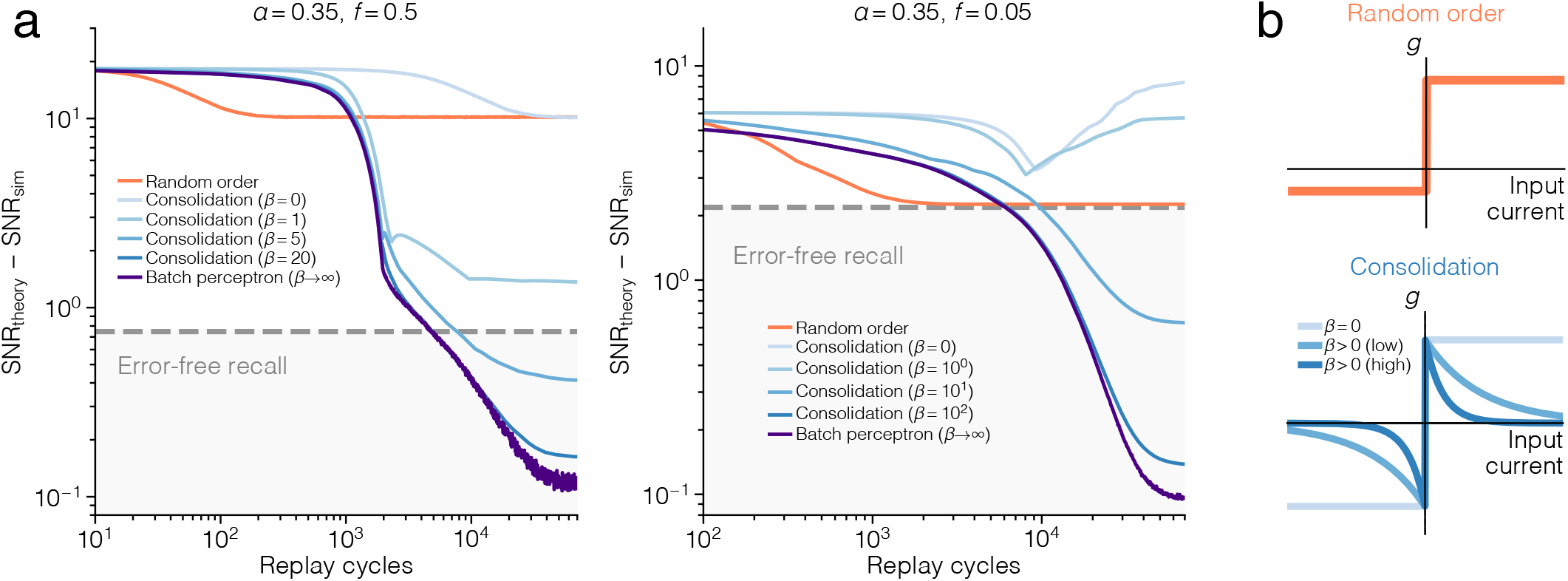
Consolidation with slow and fast gating function decay. **(a)** Difference in neural noise SNR between the theoretical optimum and the solution found by consolidating with *z* = 1 and varying, *β* values, in a single neuron (lower is better). The orange curve represents “wakeful” learning, where patterns are presented in random order and weights are updated with 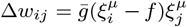. Light blue curves represent our consolidation algorithm. The dark purple curve represents our consolidation in the limit, *β* →∞, which is equivalent to the batch perceptron [81]. The dashed line indicates where the simulation crosses SNR_sim_ = 0, which is where *E* = 0 is reached. Scaling of the ordinate is arbitrary. Simulation parameters: 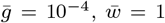, and *I*_inh_ = 8.5 for *f* = 0.5 (*I*_inh_ = 1.4 for *f* = 0.05). **(b)** Qualitative comparison of the shape of the gating function *g*, for the different variants of consolidation.

## References

[1] Frankland, P. W. & Bontempi, B. The organization of recent and remote memories. Nat. Rev. Neurosci. 6, 119–130 (2005). URL https://www.nature.com/articles/nrn1607.

[2] Wheeler, A. L. et al. Identification of a functional connectome for long-term fear memory in mice. PLoS Comput. Biol. 9, e1002853 (2013). URL 10.1371/journal.pcbi.1002853.

[3] Tonegawa, S., Morrissey, M. D. & Kitamura, T. The role of engram cells in the systems consolidation of memory. Nat. Rev. Neurosci. 19, 485–498 (2018). URL https://www.nature.com/articles/s41583-018-0031-2.

[4] Roy, D. S. et al. Brain-wide mapping reveals that engrams for a single memory are distributed across multiple brain regions. Nat. Commun. 13, 1799 (2022). URL https://www.nature.com/articles/s41467-022-29384-4.

[5] Kalisman, N., Silberberg, G. & Markram, H. The neocortical microcircuit as a tabula rasa. Proc. Natl. Acad. Sci. USA 102, 880–885 (2005). URL 10.1073/pnas.0407088102.

[6] Thomson, A. & Lamy, C. Functional maps of neocortical local circuitry. Front. Neurosci. 1 (2007). URL 10.3389/neuro.01.1.1.002.2007.

[7] Perin, R., Berger, T. K. & Markram, H. A synaptic organizing principle for cortical neuronal groups. Proc. Natl. Acad. Sci. USA 108, 5419–5424 (2011). URL https://www.pnas.org/content/108/13/5419.

[8] Khona, M. & Fiete, I. R. Attractor and integrator networks in the brain. Nat. Rev. Neurosci. 23, 744–766 (2022). URL https://www.nature.com/articles/s41583-022-00642-0.

[9] Little, W. A. The existence of persistent states in the brain. Math. Biosci. 19, 101–120 (1974). URL http://www.sciencedirect.com/science/article/pii/0025556474900315.

[10] Hopfield, J. J. Neural networks and physical systems with emergent collective computational abilities. Proc. Natl. Acad. Sci. USA 79, 2554–2558 (1982). URL https://www.pnas.org/content/79/8/2554.

[11] Gardner, E. The space of interactions in neural network models. J. Phys. A: Math. Gen. 21, 257–270 (1988). URL 10.1088%2F0305-4470%2F21%2F1%2F030.

[12] Köhler, H. M. & Widmaier, D. Sign-constrained linear learning and diluting in neural networks. J. Phys. A: Math. Gen. 24, L495–L502 (1991). URL 10.1088%2F0305-4470%2F24%2F9%2F008.

[13] Brunel, N., Hakim, V., Isope, P.Nadal, J.-P. & Barbour, B. Optimal information storage and the distribution of synaptic weights: perceptron versus Purkinje cell. Neuron 43, 745–757 (2004). URL http://www.sciencedirect.com/science/article/pii/S0896627304005288.

[14] Chapeton, J., Fares, T., LaSota, D. & Stepanyants, A. Efficient associative memory storage in cortical circuits of inhibitory and excitatory neurons. Proc. Natl. Acad. Sci. USA 109, E3614–E3622 (2012). URL http://www.pnas.org/content/109/51/E3614.

[15] Brunel, N. Is cortical connectivity optimized for storing information? Nat. Neurosci. 19, 749–755 (2016). URL https://www.nature.com/articles/nn.4286.

[16] Zhang, D., Zhang, C. & Stepanyants, A. Robust associative learning is sufficient to explain the structural and dynamical properties of local cortical circuits. J. Neurosci. 39, 6888–6904 (2019). URL https://www.jneurosci.org/content/39/35/6888.

[17] Rasch, B. & Born, J. About sleep’s role in memory. Physiol. Rev. 93, 681–766 (2013). URL 10.1152/physrev.00032.2012.

[18] Lesburguéres, E. et al. Early tagging of cortical networks is required for the formation of enduring associative memory. Science 331, 924–928 (2011). URL 10.1126/science.1196164.

[19] Kitamura, T. et al. Engrams and circuits crucial for systems consolidation of a memory. Science 356, 73–78 (2017). URL 10.1126/science.aam6808.

[20] Xu, T. et al. Rapid formation and selective stabilization of synapses for enduring motor memories. Nature 462, 915–919 (2009). URL https://www.nature.com/articles/nature08389.

[21] Chen, S. X., Kim, A. N., Peters, A. J. & Komiyama, T. Subtype-specific plasticity of inhibitory circuits in motor cortex during motor learning. Nat. Neurosci. 18, 1109–1115 (2015). URL https://www.nature.com/articles/nn.4049.

[22] Ji, D. & Wilson, M. A. Coordinated memory replay in the visual cortex and hippocampus during sleep. Nat. Neurosci. 10, 100–107 (2007). URL https://www.nature.com/articles/nn1825.

[23] Deuker, L. et al. Memory consolidation by replay of stimulus-specific neural activity. J. Neurosci. 33, 19373–19383 (2013). URL https://www.jneurosci.org/content/33/49/19373.

[24] Clawson, B. C. et al. Causal role for sleep-dependent reactivation of learning-activated sensory ensembles for fear memory consolidation. Nat. Commun. 12, 1200 (2021). URL https://www.nature.com/articles/s41467-021-21471-2.

[25] Li, W., Ma, L., Yang, G. & Gan, W.-B. REM sleep selectively prunes and maintains new synapses in development and learning. Nat. Neurosci. 20, 427–437 (2017). URL https://www.nature.com/articles/nn.4479.

[26] Zhou, Y. et al. REM sleep promotes experience-dependent dendritic spine elimination in the mouse cortex. Nat. Commun. 11, 4819 (2020). URL https://www.nature.com/articles/s41467-020-18592-5.

[27] Vyazovskiy, V. V., Cirelli, C., Pfister-Genskow, M., Faraguna, U. & Tononi, G. Molecular and electrophysiological evidence for net synaptic potentiation in wake and depression in sleep. Nat. Neurosci. 11, 200–208 (2008). URL https://www.nature.com/articles/nn2035.

[28] de Vivo, L. et al. Ultrastructural evidence for synaptic scaling across the wake/sleep cycle. Science 355, 507–510 (2017). URL https://science.sciencemag.org/content/355/6324/507.

[29] Miyamoto, D., Marshall, W., Tononi, G. & Cirelli, C. Net decrease in spine-surface GluA1-containing AMPA receptors after post-learning sleep in the adult mouse cortex. Nat. Commun. 12, 2881 (2021). URL https://www.nature.com/articles/s41467-021-23156-2.

[30] Pacheco, A. T., Bottor, J., Gao, Y. & Turrigiano, G. G. Sleep promotes downward firing rate homeostasis. Neuron 109, 1–15 (2020). URL https://www.cell.com/neuron/abstract/S0896-6273(20)30860-6.

[31] Turrigiano, G. G., Leslie, K. R., Desai, N. S., Rutherford, L. C. & Nelson, S. B. Activity-dependent scaling of quantal amplitude in neocortical neurons. Nature 391, 892–896 (1998). URL https://www.nature.com/articles/36103.

[32] Káli, S. & Dayan, P. Off-line replay maintains declarative memories in a model of hippocampal-neocortical interactions. Nat. Neurosci. 7, 286–294 (2004). URL https://www.nature.com/articles/nn1202.

[33] Tadros, T., Krishnan, G. P., Ramyaa, R. & Bazhenov, M. Sleep-like unsupervised replay reduces catastrophic forgetting in artificial neural networks. Nat. Commun. 13, 7742 (2022). URL https://www.nature.com/articles/s41467-022-34938-7.

[34] Chechik, G., Meilijson, I. & Ruppin, E. Synaptic pruning in development: a computational account. Neural Comput. 10, 1759–1777 (1998). URL 10.1162/089976698300017124.

[35] Renart, A., Song, P. & Wang, X.-J. Robust spatial working memory through homeostatic synaptic scaling in heterogeneous cortical networks. Neuron 38, 473–485 (2003). URL https://www.cell.com/neuron/abstract/S0896-6273(03)00255-1.

[36] Toyoizumi, T., Kaneko, M., Stryker, M. & Miller, K. Modeling the dynamic interaction of Hebbian and homeostatic plasticity. Neuron 84, 497–510 (2014). URL http://www.sciencedirect.com/science/article/pii/S0896627314008940.

[37] Sjöström, P. J., Turrigiano, G. G. & Nelson, S. B. Multiple forms of longterm plasticity at unitary neocortical layer 5 synapses. Neuropharmacology 52, 176–184 (2007). URL http://www.sciencedirect.com/science/article/pii/S0028390806002310.

[38] Loebel, A., Bé, J.-V. L., Richardson, M. J. E., Markram, H. & Herz, A. V. M. Matched pre- and post-synaptic changes underlie synaptic plasticity over long time scales. J. Neurosci. 33, 6257–6266 (2013). URL https://www.jneurosci.org/content/33/15/6257.

[39] Lisman, J. Glutamatergic synapses are structurally and biochemically complex because of multiple plasticity processes: long-term potentiation, long-term depression, short-term potentiation and scaling. Phil. Trans. R. Soc. B 372, 20160260 (2017). URL 10.1098/rstb.2016.0260.

[40] Cossart, R., Aronov, D. & Yuste, R. Attractor dynamics of network UP states in the neocortex. Nature 423, 283–288 (2003). URL https://www.nature.com/articles/nature01614.

[41] Nishiyama, J. & Yasuda, R. Biochemical computation for spine structural plasticity. Neuron 87, 63–75 (2015). URL http://www.sciencedirect.com/science/article/pii/S0896627315004821.

[42] Bosch, M. et al. Structural and molecular remodeling of dendritic spine sub-structures during long-term potentiation. Neuron 82, 444–459 (2014). URL http://www.sciencedirect.com/science/article/pii/S0896627314002517.

[43] Clopath, C., Ziegler, L., Vasilaki, E., Büsing, L. & Gerstner, W. Tag-trigger-consolidation: a model of early and late long-term-potentiation and depression. PLoS Comput. Biol. 4, e1000248 (2008). URL 10.1371/journal.pcbi.1000248.

[44] Redondo, R. L. & Morris, R. G. M. Making memories last: the synaptic tagging and capture hypothesis. Nat. Rev. Neurosci. 12, 17–30 (2011). URL https://www.nature.com/articles/nrn2963.

[45] Rubin, R., Abbott, L. F. & Sompolinsky, H. Balanced excitation and inhibition are required for high-capacity, noise-robust neuronal selectivity. Proc. Natl. Acad. Sci. USA 114, E9366–E9375 (2017). URL https://www.pnas.org/content/114/44/E9366.

[46] Mongillo, G., Rumpel, S. & Loewenstein, Y. Intrinsic volatility of synaptic connections — a challenge to the synaptic trace theory of memory. Curr. Opin. Neurobiol. 46, 7–13 (2017). URL http://www.sciencedirect.com/science/article/pii/S0959438817300673.

[47] Ziv, N. E. & Brenner, N. Synaptic tenacity or lack thereof: spontaneous remodeling of synapses. Trends Neurosci. 41, 89–99 (2018). URL http://www.sciencedirect.com/science/article/pii/S0166223617302370.

[48] Arellano, J. I., Benavides-Piccione, R., DeFelipe, J. & Yuste, R. Ultrastructure of dendritic spines: correlation between synaptic and spine morphologies. Front. Neurosci. 1 (2007). URL 10.3389/neuro.01.1.1.010.2007/full.

[49] Holler, S., Köstinger, G., Martin, K. A. C., Schuhknecht, G. F. P. & Stratford, K. J. Structure and function of a neocortical synapse. Nature 591, 111–116 (2021). URL https://www.nature.com/articles/s41586-020-03134-2.

[50] Hoff, P.D. Lasso, fractional norm and structured sparse estimation using a Hadamard product parametrization. Comput. Stat. Data An. 115, 186–198 (2017). URL http://www.sciencedirect.com/science/article/pii/S0167947317301469.

[51] Amid, E. & Warmuth, M. K. Winnowing with gradient descent. Proceedings of the 33rd Conference on Learning Theory, 163–182 (2020). URL http://proceedings.mlr.press/v125/amid20a.html.

[52] Loewenstein, Y., Kuras, A. & Rumpel, S. Multiplicative dynamics underlie the emergence of the log-normal distribution of spine sizes in the neocortex in vivo. J. Neurosci. 31, 9481–9488 (2011). URL http://www.jneurosci.org/content/31/26/9481.

[53] Cossell, L. et al. Functional organization of excitatory synaptic strength in primary visual cortex. Nature 518, 399–403 (2015). URL https://www.nature.com/articles/nature14182.

[54] Woloszyn, L. & Sheinberg, D. L. Effects of long-term visual experience on responses of distinct classes of single units in inferior temporal cortex. Neuron 74, 193–205 (2012). URL https://www.sciencedirect.com/science/article/pii/S0896627312001900.

[55] Fenn, K. M. & Hambrick, D. Z. Individual differences in working memory capacity predict sleep-dependent memory consolidation. J. Exp. Psychol. Gen. 141, 404 (2012). URL 10.1037/a0025268.

[56] Fenn, K. M. & Hambrick, D. Z. General intelligence predicts memory change across sleep. Psychon. Bull. Rev. 22, 791–799 (2015). URL 10.3758/s13423-014-0731-1.

[57] Ashton, J. E. & Cairney, S. A. Future-relevant memories are not selectively strengthened during sleep. PLoS ONE 16, e0258110 (2021). URL 10.1371/journal.pone.0258110.

[58] Dumay, N. Sleep not just protects memories against forgetting, it also makes them more accessible. Cortex 74, 289–296 (2016). URL https://www.sciencedirect.com/science/article/pii/S0010945215002099.

[59] Denis, D. et al. The roles of item exposure and visualization success in the consolidation of memories across wake and sleep. Learn. Mem. 27, 451–456 (2020). URL http://learnmem.cshlp.org/content/27/11/451.

[60] Jung, C. K. E. & Herms, J. Structural dynamics of dendritic spines are influenced by an environmental enrichment: an in vivo imaging study. Cereb. Cortex 24, 377–384 (2014). URL 10.1093/cercor/bhs317.

[61] Berkes, P., White, B. & Fiser, J. No evidence for active sparsification in the visual cortex. Advances in Neural Information Processing Systems 22 (2009). URL https://proceedings.neurips.cc/paper/2009/hash/2b24d495052a8ce66358eb576b8912c8-Abstract.html.

[62] Alemi, A., Baldassi, C., Brunel, N. & Zecchina, R. A three-threshold learning rule approaches the maximal capacity of recurrent neural networks. PLoS Comput. Biol. 11, e1004439 (2015). URL 10.1371/journal.pcbi.1004439.

[63] Kaufman, M., Corner, M. A. & Ziv, N. E. Long-term relationships between cholinergic tone, synchronous bursting and synaptic remodeling. PLoS ONE 7, e40980 (2012). URL 10.1371/journal.pone.0040980.

[64] Hazan, L. & Ziv, N. E. Activity dependent and independent determinants of synaptic size diversity. J. Neurosci. 40, 2828–2848 (2020). URL https://www.jneurosci.org/content/40/14/2828.

[65] Fisher-Lavie, A. & Ziv, N. E. Matching dynamics of presynaptic and postsynaptic scaflolds. J. Neurosci. 33, 13094–13100 (2013). URL https://www.jneurosci.org/content/33/32/13094.

[66] Gala, R. et al. Computer assisted detection of axonal bouton structural plasticity in in vivo time-lapse images. eLife 6, e29315 (2017). URL 10.7554/eLife.29315.

[67] Ishii, K. et al. In vivo volume dynamics of dendritic spines in the neocortex of wild-type and Fmr1 KO mice. eNeuro 5, e0282.#x2013;18.2018 (2018). URL http://www.eneuro.org/content/5/5/ENEURO.0282-18.2018.

[68] Steffens, H. et al. Stable but not rigid: chronic in vivo STED nanoscopy reveals extensive remodeling of spines, indicating multiple drivers of plasticity. Sci. Adv. 7, eabf2806 (2021). URL 10.1126/sciadv.abf2806.

[69] Wegner, W., Steffens, H., Gregor, C., Wolf, F. & Willig, K. I. Environmental enrichment enhances patterning and remodeling of synaptic nanoarchitecture as revealed by STED nanoscopy. eLife 11, e73603 (2022). URL 10.7554/eLife.73603.

[70] Morrison, A., Aertsen, A. & Diesmann, M. Spike-timing-dependent plasticity in balanced random networks. Neural Comput. 19, 1437–1467 (2007). URL 10.1162/neco.2007.19.6.1437.

[71] Miller, K. D. & MacKay, D. J. C. The role of constraints in Hebbian learning. Neural Comput. 6, 100–126 (1994). URL 10.1162/neco.1994.6.1.100.

[72] Sacramento, J., Wichert, A. & van Rossum, M. C. W. Energy efficient sparse connectivity from imbalanced synaptic plasticity rules. PLoS Comput. Biol. 11, e1004265 (2015). URL 10.1371/journal.pcbi.1004265.

[73] Shouval, H. Z. Clusters of interacting receptors can stabilize synaptic efficacies. Proc. Natl. Acad. Sci. U.S.A. 102, 14440–14445 (2005). URL 10.1073/pnas.0506934102.

[74] Triesch, J., Vo, A. D. & Hafner, A.-S. Competition for synaptic building blocks shapes synaptic plasticity. eLife 7, e37836 (2018). URL 10.7554/eLife.37836.

[75] Benna, M. K. & Fusi, S. Computational principles of synaptic memory consolidation. Nat. Neurosci. 19, 1697–1706 (2016). URL https://www.nature.com/articles/nn.4401.

[76] Li, H. L. & van Rossum, M. C. W. Energy efficient synaptic plasticity. eLife 9, e50804 (2020). URL 10.7554/eLife.50804.

[77] Euston, D. R., Tatsuno, M. & McNaughton, B. L. Fast-forward playback of recent memory sequences in prefrontal cortex during sleep. Science 318, 1147–1150 (2007). URL 10.1126/science.1148979.

[78] Crick, F. & Mitchison, G. The function of dream sleep. Nature 304, 111–114 (1983). URL https://www.nature.com/articles/304111a0.

[79] Hopfield, J. J., Feinstein, D. I. & Palmer, R. G. ‘Unlearning’ has a stabilizing effect in collective memories. Nature 304, 158–159 (1983). URL https://www.nature.com/articles/304158a0.

[80] Nacson, M. S. et al. Convergence of gradient descent on separable data. Proceedings of the 22nd International Conference on Artificial Intelligence and Statistics, 3420–3428 (2019). URL http://proceedings.mlr.press/v89/nacson19b.html. ISSN: 2640-3498.

[81] Krauth, W. & Mezard, M. Learning algorithms with optimal stability in neural networks. J. Phys. A: Math. Gen. 20, L745–L752 (1987). URL 10.1088%2F0305-4470%2F20%2F11%2F013.

[82] Bouten, M., Engel, A., Komoda, A. & Serneels, R. Quenched versus annealed dilution in neural networks. J. Phys. A: Math. Gen. 23, 4643 (1990). URL 10.1088/0305-4470/23/20/025.

[83] Yau, H. W. Phase space techniques in neural network models. PhD thesis, University of Edinburgh, Edinburgh, Scotland (1992). URL https://era.ed.ac.uk/handle/1842/14713.

[84] Tsodyks, M. V. & Feigel’man, M. V. The enhanced storage capacity in neural networks with low activity level. Europhys. Lett. 6, 101–105 (1988). URL 10.1209%2F0295-5075%2F6%2F2%2F002.

[85] Rolls, E. T. & Tovee, M. J. Sparseness of the neuronal representation of stimuli in the primate temporal visual cortex. J. Neurophysiol. 73, 713–726 (1995). URL 10.1152/jn.1995.73.2.713.

[86] Vinje, W. E. & Gallant, J. L. Sparse coding and decorrelation in primary visual cortex during natural vision. Science 287, 1273–1276 (2000). URL 10.1126/science.287.5456.1273.

[87] Wickens, T. D. Elementary signal detection theory. Oxford University Press (2002).

[88] Cortes, C. & Vapnik, V. Support-vector networks. Mach. Learn. 20, 273–297 (1995). URL 10.1007/BF00994018.

[89] Mangasarian, O. L. Arbitrary-norm separating plane. Oper. Res. Lett. 24, 15–23 (1999). URL https://www.sciencedirect.com/science/article/pii/S0167637798000492.

[90] Oja, E. Simplified neuron model as a principal component analyzer. J. Math. Biology 15, 267–273 (1982). URL 10.1007/BF00275687.

[91] Chechik, G., Meilijson, I. & Ruppin, E. Neuronal regulation: a mechanism for synaptic pruning during brain maturation. Neural Comput. 11, 2061–2080 (1999). URL 10.1162/089976699300016089.

[92] Zenke, F. & Ganguli, S. SuperSpike: supervised learning in multilayer spiking neural networks. Neural Comput. 30, 1514–1541 (2018). URL 10.1162/necoa01086.

[93] Amit, D. J., Campbell, C. & Wong, K. Y. M. The interaction space of neural networks with sign-constrained synapses. J. Phys. A: Math. Gen. 22, 4687–4693 (1989). URL 10.1088%2F0305-4470%2F22%2F21%2F030.

